# BRAF inhibitors reprogram cancer-associated fibroblasts to drive matrix remodeling and therapeutic escape in melanoma

**DOI:** 10.1101/2021.02.18.431841

**Authors:** Tianyi Liu, Linli Zhou, Yao Xiao, Thomas Andl, Yuhang Zhang

## Abstract

The tumor stroma and its cellular components are known to play an important role in tumor resilience towards treatment. Here, we report a novel resistance mechanism in melanoma that is elicited by BRAF inhibitors (BRAFi)-induced non-canonical nuclear β-catenin signaling in cancer-associated fibroblasts (CAFs). Our study reveals that BRAFi leads to an expanded CAF population with increased β-catenin nuclear accumulation and enhances their biological properties. This CAF subpopulation is essential for melanoma cells to grow and resist to BRAFi/MEK inhibitors (MEKi). Mechanistically, BRAFi induces BRAF and CRAF dimerization and subsequently the activation of ERK signaling in CAFs, leading to the inactivation of the β-catenin destruction complex. By RNA-Seq, periostin (POSTN) is identified as a major downstream effector of β catenin in CAFs and can compensate for the loss of β-catenin in CAFs in conferring melanoma cell BRAFi/MEKi resistance. Moreover, POSTN reactivates the ERK pathway, which is inhibited by BRAFi/MEKi, in melanoma cells through PI3K/AKT signaling. Our data underscore the roles of BRAFi-induced CAF reprogramming in matrix remodeling and therapeutic escape of melanoma cells and reveal POSTN as an important matrix target to eliminate BRAFi/MEKi resistance in melanoma.

## Main

Cutaneous melanoma is the deadliest type of skin cancer and one of the leading causes of cancer-related mortality due to its metastatic potential^1^. Over the past decade, an improved understanding of the mutational landscape in melanoma has led to the identification of BRAF V600 mutations in about 60% of patients^2^. The discovery spurred the development of specific inhibitors for the pathways that are affected by BRAF mutations^3^, namely BRAFi. However, therapeutic outcomes are disappointing for many melanoma patients due to the occurrence of drug resistance^4^. Because many resistance mechanisms involve mitogen-activated protein kinase (MAPK) reactivation, BRAFi and MEKi combination therapies have emerged as major treatment options^5^. Unfortunately, despite the fact that combined BRAFi and MEKi therapies have shown improved response rates and prolonged progression-free survival, melanoma resistance and recurrence still frequently happen in most patients after six months^6^.

Resistance to targeted therapies is a persistent hurdle in malignant melanoma treatment and management and can occur through multiple mechanisms. However, current studies of melanoma drug resistance mainly focus on intrinsic molecular alternations, such as mitogen-activated protein kinase (MAPK) reactivation, and extrinsic resistance mechanisms are underexplored. Tumors are complex tissues consisting of mixed populations of cancer cells and stromal cells embedded in the dense and stiff extracellular matrix (ECM)^7^. These heterogeneous cell populations produce a myriad of cell-cell and cell-ECM interactions, contributing to tumor cell adaptation and, eventually, tumor recurrence and end-stage disease in most patients^8^. An increased number of studies reported that treatment-induced changes could occur in multiple stromal cell compartments and lead to the development of drug resistance phenotypes and reduced efficacy of therapies^9–13^. It was also reported that treatment-relapsed melanomas converge on a shared resistance phenotype characterized by remodeled ECM and increased tumor stiffness, altering treatment responses in melanoma cells^14^. Therefore, in the era of combinatorial therapy, the tumor stroma and its cellular components cannot be ignored or underestimated in order to quell tumor resilience towards treatment and to improve therapeutic outcomes for melanoma patients.

One notorious key component of the tumor stroma is cancer-associated fibroblasts (CAFs)^15^. Although many other stromal cell types and tumor cells can produce ECM proteins, CAFs appear to be the major player in the stroma that synthesizes, secrets, assembles, and modifies the ECM composition, organization, and stiffness^16–18^. Particularly, CAFs are known to produce ECM proteins, including fibrillar collagens and fibronectin, ECM-degrading enzymes, such as matrix metalloproteinases (MMPs), and collagen cross-linking enzymes, such as members of the lysyl oxidase (LOX) family. Therefore, CAFs could be a rich and complex source for melanoma cells to rely on in case of a therapeutic attack. However, CAFs are not a monolithic cell population and include a dynamic collection of fibroblast subsets with different characteristics and functions^19–21^. For example, the Tuveson group identified two distinct CAF subpopulations in the pancreatic tumor microenvironment, a myofibroblast-like CAF population, termed “myCAFs,” and an inflammatory CAF population, termed “iCAFs”^22^. As such, a better understanding of heterogeneous CAF populations in the ever-changing stromal microenvironment, such as the presence of administrated therapeutic agents, is urgently needed in order to solve the mystery of melanoma drug resistance.

We discovered that BRAFi leads to the expansion of a subset of CAFs with increased nuclear β-catenin accumulation. Previously, we reported that β-catenin, a molecule driving normal and pathological responses in fibroblasts during wound healing, fibrosis, and keloid pathogenesis^23^, is essential for the biological properties of dermal fibroblasts^24^. We demonstrated that targeted β-catenin ablation in CAFs led to suppressed melanoma growth *in vitro* and *in vivo*^25^. The findings suggest that β-catenin could be a master regulator of a subset of readapted CAFs in melanoma under the siege of therapeutic agents.

In this study, we have investigated how BRAFi stimulates CAFs through nuclear β-catenin activation to remodel the ECM microenvironment, eliciting alternative signaling in BRAF-mutant melanoma cells to bypass BRAFi/MEKi inhibition. We found that BRAFi leads to an enhanced phenotype in CAFs by inducing BRAF and CRAF dimerization and increased nuclear β-catenin accumulation. By RNA-Seq, we identified matricellular protein periostin (POSTN) as a downstream effector of β catenin in CAFs. POSTN is uniquely expressed in CAFs, and BRAFi can specifically upregulate POSTN production in CAFs but not in melanoma cells. Recombinant POSTN can compensate for the loss of β-catenin in CAFs and contribute to melanoma cell BRAFi/MEKi resistance. Importantly, our study reveals that POSTN activates PI3K/AKT signaling in melanoma cells, which reactivates the ERK signaling pathway that is inhibited by BRAFi and MEKi. Together, our data reveal a novel BRAFi-stimulated CAF-mediated targeted drug resistance pathway in melanoma and POSTN as an exciting therapeutic candidate to overcome melanoma resistance to BRAFi/MEKi.

## Results

### A subset of CAFs with increased nuclear β-catenin accumulation is expanded in BRAFi/MEKi-treated melanoma

CAFs are known to be a notorious ECM-remodeling machine in the tumor stroma^26^. To understand the roles of CAFs in melanoma resistance to BRAFi and MEKi, we performed a paired-analysis of melanoma tissue samples collected from patients carrying BRAF V600-mutated melanomas before and after BRAFi/MEKi therapy (Table 1). CAFs in melanoma were identified using a known CAF marker, alpha-smooth muscle actin (α-SMA)^27^. Notably, the total number of α-SMA-positive (α-SMA+) CAFs was increased in melanomas isolated from the same patients after BRAFi/MEKi treatment (Fig. 1e). Most importantly, the numbers of α-SMA+ CAFs expressing nuclear β-catenin were also increased after BRAFi/MEKi treatment (Fig. 1b, d) compared with the ones before BRAFi/MEKi treatment (Fig. 1a, c). The percentage of α-SMA+ cells expressing nuclear β-catenin was increased from 17 ± 2% before treatment to 41 ± 2% after BRAFi/MEKi treatment (Fig. 1f). The data were validated using ten human melanoma samples collected from the patients without udergoing BRAFi/MEKi treatment (Table 2), six samples collected from the patients who received BRAFi treatment (Table 3, patient number 19-24), and six samples collected from the patients who received BRAFi/MEKi treatment (Table 3, patient number 25-30).

**Fig. 1.**
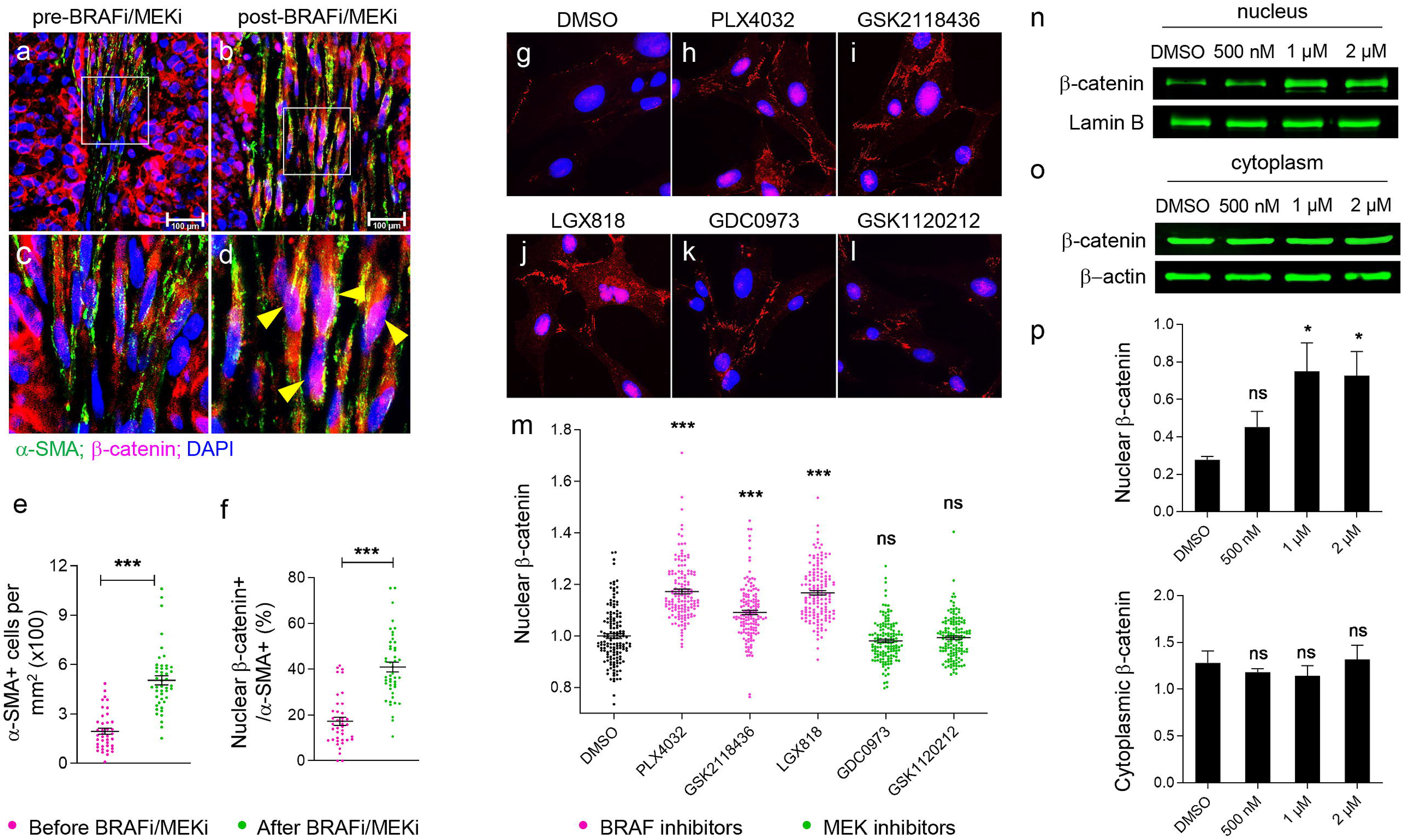
BRAFi drives increased nuclear β-catenin in CAFs. **a-d**. Human melanoma sections (collected from eight melanoma patients before (**a**) and after (**b**) BRAFi/MEKi therapy) were co-immunostained for α-SMA and β-catenin expression using an anti-α-SMA antibody (green) and an anti-β-catenin antibody (red). The nuclei were stained blue with DAPI. **c** and **d** are close-up pictures of the area circled by white boxes in **a** and **b**, respectively. Yellow triangles in **d** indicate α-SMA+ fibroblasts expressing nuclear β-catenin. Scale bar: 100 µm. **e**. Scattered plot represents the numbers of α-SMA+ cells per mm^2^ counted in pre- and post-treated melanoma samples. **f**. Percentage of α-SMA+ cells expressing nuclear β-catenin in pre- and post-treated melanoma samples. Each data point represents the quantification result of one picture. At least five pictures were taken per tumor section. n ≥ 40. **g-l**. Immunostaining of β-catenin in M50 treated with DMSO or indicated inhibitors for 72 hours. **m**. Quantification and comparison of relative fluorescence intensities of nuclear β catenin in M50 treated with DMSO or indicated BRAFi/ MEKi for 72 hours. Fluorescence intensity of nuclear β-catenin in DMSO-treated M50 was used as the control and set at one for comparison. Each data point represents the relative fluorescent intensities of nuclear β-catenin in one treated cell. n=145. **n**. Nuclear and cytoplasmic fractions of M50 treated with DMSO or different concentrations of PLX4032 were extracted for β-catenin Western blotting. **o**. Fluorescence intensities of nuclear and cytoplasmic β-catenin bands in Western blotting were quantified and compared from three independent experiments. n=3. For all graphs, data are represented as mean ± SEM. * *P* ≤ 0.05. ** *P* ≤ 0.01. *** *P* ≤ 0.001. ns: not significant.

**Table 1.** Paired human melanoma samples collected from the same melanoma patients before and after BRAFi/MEKi therapy.

| Number | Gender | Subtype | Sample ID |  | Therapy |
| --- | --- | --- | --- | --- | --- |
|  |  |  | Pre-resistance sample | Post resistance sample |  |
| 1 | M | SSM | S09-008757 A7 | S17-002953 B1 | BRAFi/MEKi combo |
| 2 | M | NOS | S11-006710 A1 | S14-001866 A1 | BRAFi/MEKi combo |
| 3 | F | Nodular | S08-017787 D3 | S12-020204 A1 | BRAFi then BRAF/MEKi |
| 4 | F | Unknown primary | S14-005069 A1 | S14-018628 B1 | BRAFi/MEKi combo |
| 5 | M | SSM | S10-009993 A2 | S12-006873 A5 | BRAFi then BRAF/MEKi |
| 6 | M | SSM | S15-007219 A1 | S15-018894 A1 | BRAFi/MEKi combo |
| 7 | M | SSM | S13-007345 C15 | S18-13781 A2 | BRAFi then BRAF/MEKi |
| 8 | F | SSM | S14-007782 B4 | S18-22658 A3 | BRAFi/MEKi combo |

**Table 2.**
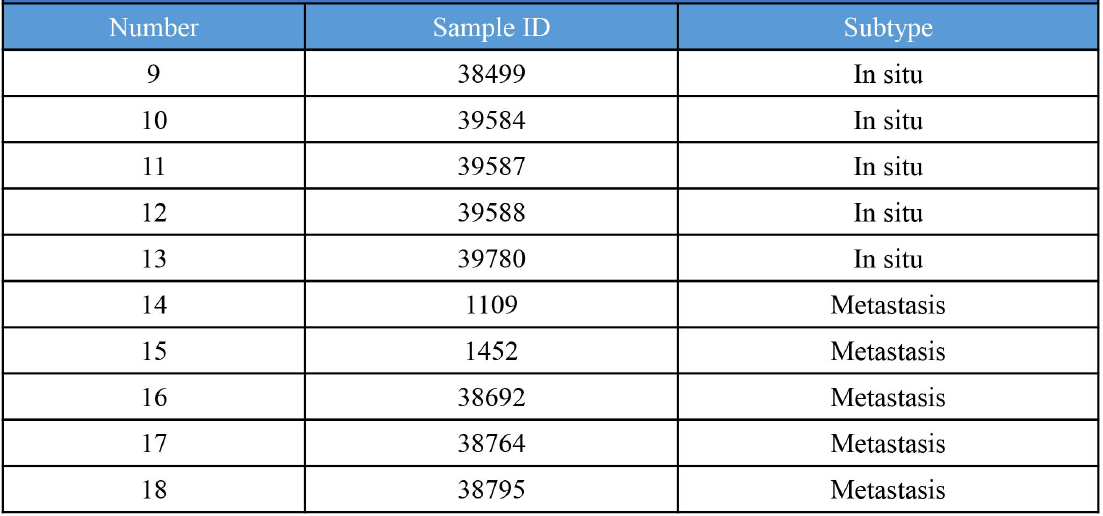
Human melanoma samples collected from the patients without BRAFi/MEKi treatment.

**Table 3.** Human melanoma samples collected from the patients who received BRAFi or BRAFi/MEKi treatment.

| Number | Sample ID | Gender | Subtype | Therapy |
| --- | --- | --- | --- | --- |
| 19 | DC1 | F | Skin-Chest | BRAFi |
| 20 | DC3 | M | Lymph Node-Groin | BRAFi |
| 21 | DC5 | F | Skin-Lower leg (in transit) | BRAFi |
| 22 | MAL01124 | F | Skin-foot | BRAFi |
| 23 | MAL01124 | F | soft tissue-lower leg | BRAFi |
| 24 | THO00841 | M | Lung | BRAFi |
| 25 | S13-002767 A1 | M | SSM | BRAFi/MEKi combo |
| 26 | S12-16290 B2 | F | NOS | BRAFi/MEKi combo |
| 27 | S13-008996 A3 | F | Unknown primary | BRAFi then BRAF/MEKi |
| 28 | S15-016538-A1 | F | Nodular | BRAFi/MEKi combo |
| 29 | DC4 | F | Skin-Groin (met) | BRAFi/MEKi combo |
| 30 | MAL000267 | M | Lymph Node | BRAFi/MEKi combo |

### BRAFi drives increased nuclear **β**-catenin in CAFs

To understand whether increased nuclear β-catenin in CAFs is driven by BRAFi or MEKi, we obtained six CAF cell lines isolated from surgically excised human melanoma tissues from two different sources (M27, M28, M50, M56, M77, and M80). We confirmed that they express increased levels of known CAF markers, including α-SMA, fibroblast activation protein (FAP)^28^, and fibroblast-specific protein-1 (FSP-1, also known as S100A4)^27^ as compared with normal human dermal fibroblast (Fig. S1a-c). Quantification analysis revealed that M27 and M50 express the highest levels of α-SMA, FAP, and FSP-1 among all CAF cell lines (Table 4). In addition, all CAF cell lines exhibit pronounced filamentous actin (F-actin) expression (Fig. S1d), which is known to be involved in cell migration and division^29^. As shown by the scratch assay, which is a direct reflection of cell contractility and migration, all six CAF cell lines exhibit increased migratory abilities than normal human dermal fibroblasts (Fig. S1e). Collagen gel contraction assays were performed to measure the ability of the fibroblasts to remodel the ECM. Six CAF cell lines showed enhanced but variable ability to contract the gel, among which M27 and M50 showed the strongest abilities to contract collagen gels (Fig. S1f).

**Table 4.** Comparison of the expression levels of indicated proteins, migration, and gel contraction between six CAF lines and normal fibroblasts. To assess the fluorescence intensities of expressed molecules in fibroblasts shown in Fig. S1, the slides were photographed using a Cytation 1 multi-mode cell imager, and fluorescence intensity in each cell was measured using GEN 5.3 software (BioTek, Winooski, VT) (n≥75). The highest fluorescence intensity was set to five (indicated by “+++++”) to establish a scoring scale for each immunofluorescent staining. Similar approaches were used to create a scoring scale for gel contraction (n=3) and *in vitro* cell migration assays (n=4). M27 and M50 showed the highest scores among all tested CAF lines and normal fibroblasts.

|  | a-SMA | FAP | FSP-1 | F-actin | Gel contraction | Migration |
| --- | --- | --- | --- | --- | --- | --- |
| NF | + | + | + | + | ++ | +++ |
| M27 | +++++ | +++++ | ++++ | ++++ | +++++ | ++++ |
| M28 | ++++ | +++ | ++ | ++++ | +++ | ++++ |
| M50 | +++++ | +++++ | +++++ | +++++ | +++++ | +++++ |
| M56 | ++++ | +++ | ++ | +++ | +++ | ++++ |
| M77 | +++ | ++++ | ++++ | ++++ | ++ | +++++ |
| M80 | ++ | +++++ | ++++ | ++++ | ++ | ++++ |

Next, we investigated whether BRAFi, including PLX4032 (vemurafnib), GSK2118436 (dabrafenib), and LGX818 (encorafenib), or MEKi, including GDC0973 (cobimetinib), and GSK1120212 (Trametinib), can lead to increased nuclear β-catenin in CAFs. Interestingly, all three BRAF inhibitors led to increased nuclear β-catenin in M50 (Fig. 1h-j) while the nuclear β-catenin levels in MEKi-treated M50 remained unchanged (Fig. 1k-l) as compared with control DMSO-treated cells (Fig. 1g), suggesting that increased nuclear β-catenin in CAFs is driven by BRAFi but not MEKi (Fig. 1m). The nuclear and cytoplasmic fractions of M50 treated with either DMSO or different concentrations of PLX4032 (500 nM, 1 µM, and 2 µM) were extracted for Western blotting analysis. As shown in Fig. 1p, PLX4032-treated M50 show increased amounts of nuclear β-catenin in a concentration-dependent manner (Fig. 1n), while the levels of cytoplasmic β-catenin in M50 remain unchanged under PLX4032 treatment (Fig. 1o).

### PLX4032 stimulates the biological properties of CAFs

Surprisingly, we discovered that PLX4032 alters the biological properties of CAFs. Seventy-two hours after PLX4032 treatment, not only there was an increase in nuclear β-catenin (Fig. 2a), M50 exhibited a more pronounced F-actin expression (Fig. 2b) and increased levels of focal adhesion protein paxillin (Fig. 2c) and phosphorylated myosin light chain 2 (p-MLC2) (Fig. 2d). We next asked whether PLX4032 can enhance the biological functions of CAFs. Surprisingly, although M50 treated with DMSO (vehicle) already contracted collagen gel (37.59 % ± 2.15), PLX4032 further increased the gel contracting ability of M50 (61.42% ± 2.92) (Fig. 2e). More significantly, confocal reflection microscopy (CRM) revealed increased fiber connectivity in the gel embedded with PLX4032-treated M50 (9.82 ± 1.23) (Fig. 2f). Connectivity measured by ImageJ denotes the extent of line connections in a network, indicating the complexity of collagen fibers in the gel. Furthermore, the migratory ability of M50 was significantly increased by PLX4032 (87.76 ± 1.64%) comparing to M50 treated with DMSO (68.02 ± 1.49%) (Fig. 2g). The stiffness of the gels embedded with M50 increased substantially from 1038 ± 70.21 Pa to 5863 ± 73.52 Pa upon PLX4032 treatment (Fig. 2h). The data collectively reveal that PLX4032 enhances cytoskeletal dynamics in CAFs and their matrix remodeling ability.

**Fig. 2.**
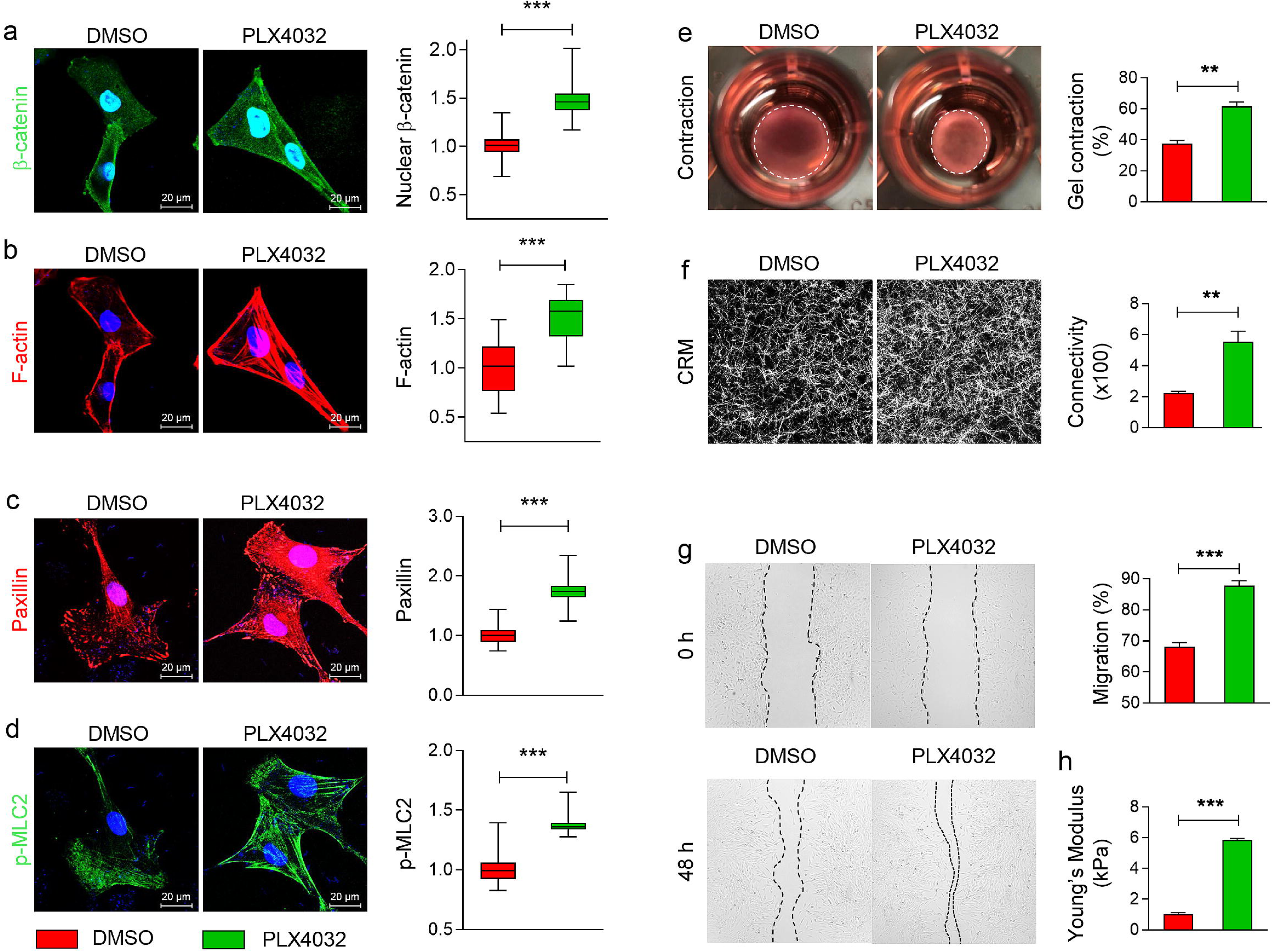
PLX4032 stimulates the biological properties of CAFs. M50 was treated with either DMSO or PLX4032 for 72 hours for evaluation. **a**-**d**. β-catenin, F-actin, paxillin, and p-MLC2 immunostaining of DMSO-or PLX4032-treated M50. The nuclei were stained blue with DAPI. All immunostainings were repeated a minimum of three times. Scale bar: 20 μm. Fluorescence intensities of nuclear β catenin, F-actin, paxillin and p-MLC2 were normalized by the mean intensities of protein expression in DMSO-treated M50 and presented in a box and whisker plot. n≥39. **e**. Representative images of collagen gels contracted by M50 treated with DMSO or PLX4032 after 72 hours. Bar graph shows percentages of gel contraction from three independent experiments. **f**. Representative CRM images of the gels embedded with M50 treated with DMSO or PLX4032 after 72 hours. The graph shows the connectivity of collagen fibers in gels remodeled by M50 from three independent experiments. **g**. Images show M50 migration under DMSO or PLX4032 treatment at 0 and the 48th hour in an *in vitro* scratch assay. Black short dash lines depict the frontlines of M50 at indicated time points. Bar graph shows migration percentages of M50 at the 48th hour under DMSO or PLX4032 treatment from four independent experiments. **h**. The graph shows Young’s modulus of collagen gels, which were remodeled by M50 treated with DMSO or PLX4032 for 72 hours, measured by AFM. n≥10. In all bar graphs, data represent mean ± SEM. ** P ≤ 0.01. *** P ≤ 0.001. ns: not significant.

### **β**-catenin depletion eliminates the ability of CAFs to respond to PLX4032 stimulation

To understand whether β-catenin is essential for CAFs to respond to BRAFi, β-catenin expression in M27 and M50 was silenced using a doxycycline-inducible short hairpin RNA (shRNA) as we have reported before^24^ and tagged with green fluorescent protein (GFP). M50 and M27 cells carrying β-catenin shRNA are designated as bcat-GFP/M50 and bcat-GFP/M27. Control GFP/M50 and GFP/M27 were generated by transducing the cells with a GFP-tagged non-targeting lentiviral shRNA. Western blotting demonstrated that the expression of β was ablated efficiently in bcat-GFP/M50 (Fig. S2a-b).

Depleting β-catenin reduced the proliferation of M50 cells as indicated by cell number counting (Fig. S2c) and the MTT assay (Fig. S2d). Suppressed M50 growth was mainly caused by decreased cell proliferation but not increased cell apoptosis. As shown in Fig. S2g, 90 ± 1% of GFP/M50 cells (Fig. S2e) were Ki67-positive (Ki67+) while only 72 ± 2% of bcat-GFP/M50 cells (Fig. S2f) were Ki67+. No significant difference in the numbers of TUNEL-positive (TUNEL+) cells (Fig. S2j) was detected between GFP/M50 (Fig. S2h) and bcat-GFP/M50 (Fig. S2i). More importantly, β-catenin deficiency weakened the ability of M50 cells to contract the collagen gel (Fig. S2k-l). Similarly, the density of collagen fibers in the gel embedded with bcat-GFP/M50 was lower than that of the gel embedded with GFP/M50 (Fig. S2m). Quantification in Fig. S2n shows that the connectivity of collagen fibers in gel embedded with bcat-GFP/M50 was significantly (0.54 ± 0.07) less than that of the gel embedded with GFP/M50 (4.64 ± 0.23). β-catenin deficiency also reduced the abilities of M50 to migrate as shown by the scratch assay (Fig. S2o, p). The stiffness of the gel embedded with bcat-GFP/M50 was 122.0 ± 9.25 Pa compared with 1007 ± 32.21 Pa for the gel embedded with GFP/M50 (Fig. S2q).

Interestingly, upon β-catenin depletion (Fig. 3a), M50 cells not only exhibited reduced levels of F-actin (Fig. 3b), paxillin (Fig. 3c), and p-MLC2 (Fig. 3d) and no longer responded to PLX4032. Furthermore, PLX4032 failed to enhance the gel contracting ability of β-catenin-deficient M50 cells (Fig. 3e). CRM did not detect increased fiber connectivity in the gel embedded with PLX4032-treated β-catenin-deficient M50 (Fig. 3f). The migratory ability of β-catenin-deficient M50 cells also remained unchanged upon PLX4032 treatment comparing to M50 cells treated with PLX4032 (Fig. 3g). The data demonstrate that β-catenin activity is essential for CAFs to respond to PLX4032. The data suggest that depleting β-catenin could be an effective way to eliminate this unique subset of BRAFi-stimulated CAFs.

**Fig. 3.**
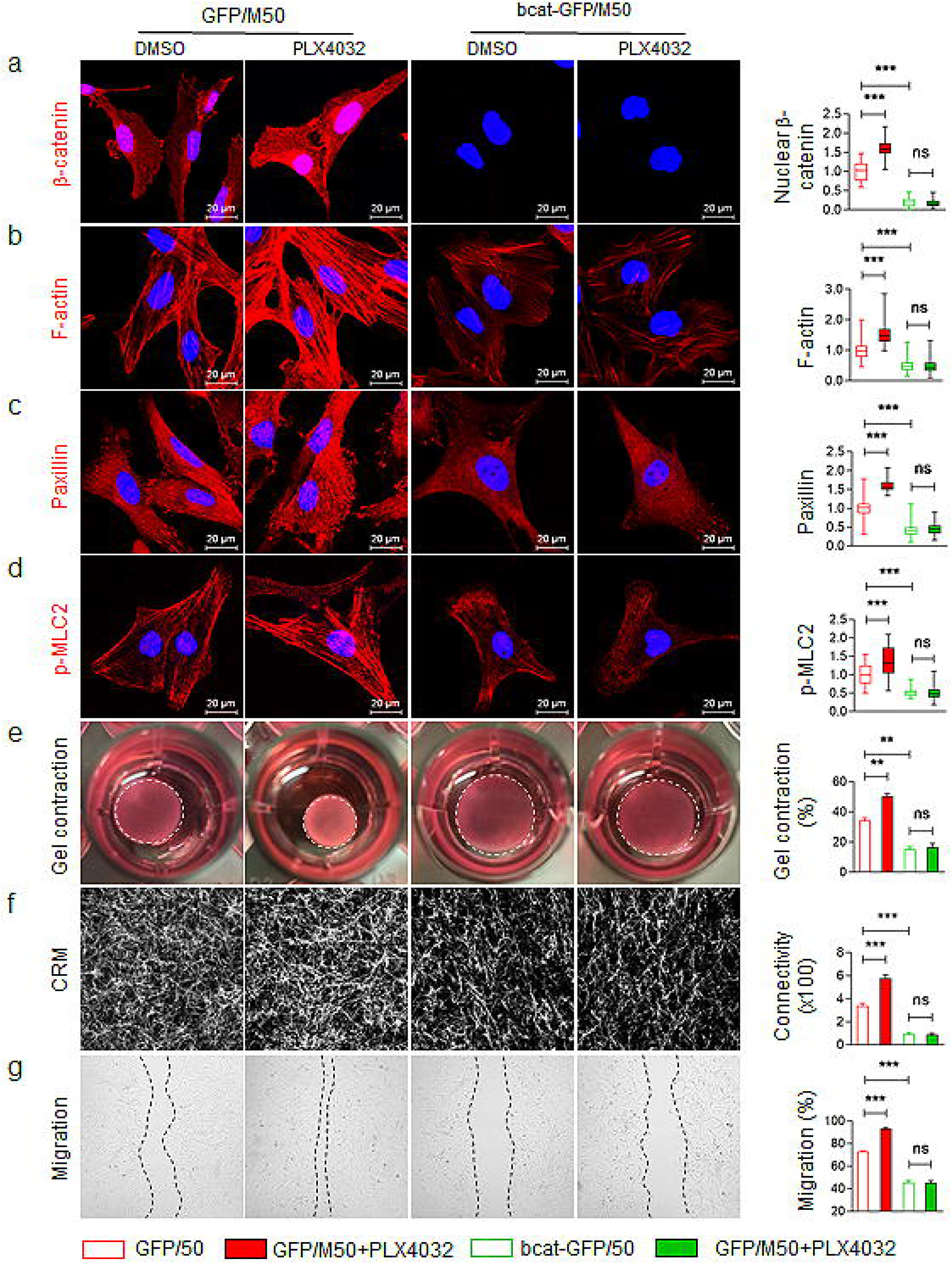
β-catenin depletion eliminates the ability of CAFs to respond to PLX4032 stimulation. **a-d**. GFP/M50 and bcat-GFP/M50 were treated with either DMSO or PLX4032 for 72 hours for β catenin (**a**), F-actin (**b**), paxillin (**c**) and p-MLC2 (**d**) immunostaining. The nuclei were stained blue with DAPI. Scale bar: 20 μm. Fluorescence intensities of nuclear β-catenin, F-actin, paxillin and p-MLC2 were normalized by the mean intensities of protein expression in DMSO-treated M50 and presented in a box and whisker plot. n=100. **e**. Representative images of collagen gels contracted by GFP/M50 and bcat-GFP/M50 treated with DMSO or PLX4032 after 72 hours. Percentages of gel contraction were measured and shown as mean ± SEM in bar graph. n=3. **f**. Representative CRM images of the gels embedded with GFP/M50 and bcat-GFP/M50 treated with DMSO or PLX4032 after 72 hours. The graph shows the connectivity of fibers in collagen gels remodeled by M50 (mean ± SEM). n=7. **g**. Images show GFP/M50 and bcat-GFP/M50 migration under DMSO or PLX4032 treatment after 48 hours in an *in vitro* scratch assay. Black short dash lines depict the frontlines of GFP/M50 and bcat-GFP/M50 at the end point. The graph shows migration percentage of GFP/M50 or bcat-GFP/M50 and expressed as mean ± SEM. n=9. For all bar graphs, data are represented as mean ± SEM. ** *P* ≤ 0.01. *** *P* ≤ 0.001. ns: not significant.

### BRAFi-stimulated CAFs contribute to melanoma cell resistance to BRAFi and MEKi

To understand whether inhibiting this subset of CAFs can suppress BRAF-mutant melanoma cell resistance to BRAFi and MEKi, we designed a multicolor 3D co-culture system consisting of human melanoma cells and CAFs. BRAF-mutant melanoma cell lines, including A375 and SK-Mel-24, were labeled with red fluorescent protein (RFP) using a non-targeting shRNA carrying an RFP-encoding gene. Mitomycin-treated CAFs (designated as mito-GFP/M27 and mito-GFP/M50) were used as non-functional CAF controls.

As shown in Fig. 4a, b, after a 72-hour co-culture, there were 29 ± 4 % more A375 cells in A375 + GFP/M50 spheroids compared with A375 + mito-GFP/M50 spheroids. However, no significant difference in A375 numbers was observed between A375 + bcat-GFP/M50 and A375 + mito-GFP/M50, indicating β-catenin depletion deprived M50 cells of their ability to support A375 growth. Because BRAF V600E mutation leads to abnormal activation of the MAPK/ERK signaling pathway^30^, we examined the level of phospho-ERK (p-ERK) in A375 in 3D co-cultured spheroids. M50 in the spheroids were identified using a fibroblast-specific antibody, TE7^31^. As shown in Fig. 4c, while only 42 ± 2% A375 expressed p-ERK when co-cultured with mito-GFP/M50, 56 ± 2% of A375 co-cultured with GFP/M50 expressed p-ERK. β-catenin ablation in M50 cells had negative effects on ERK signaling in A375 (39 ± 1%).

**Fig. 4.**
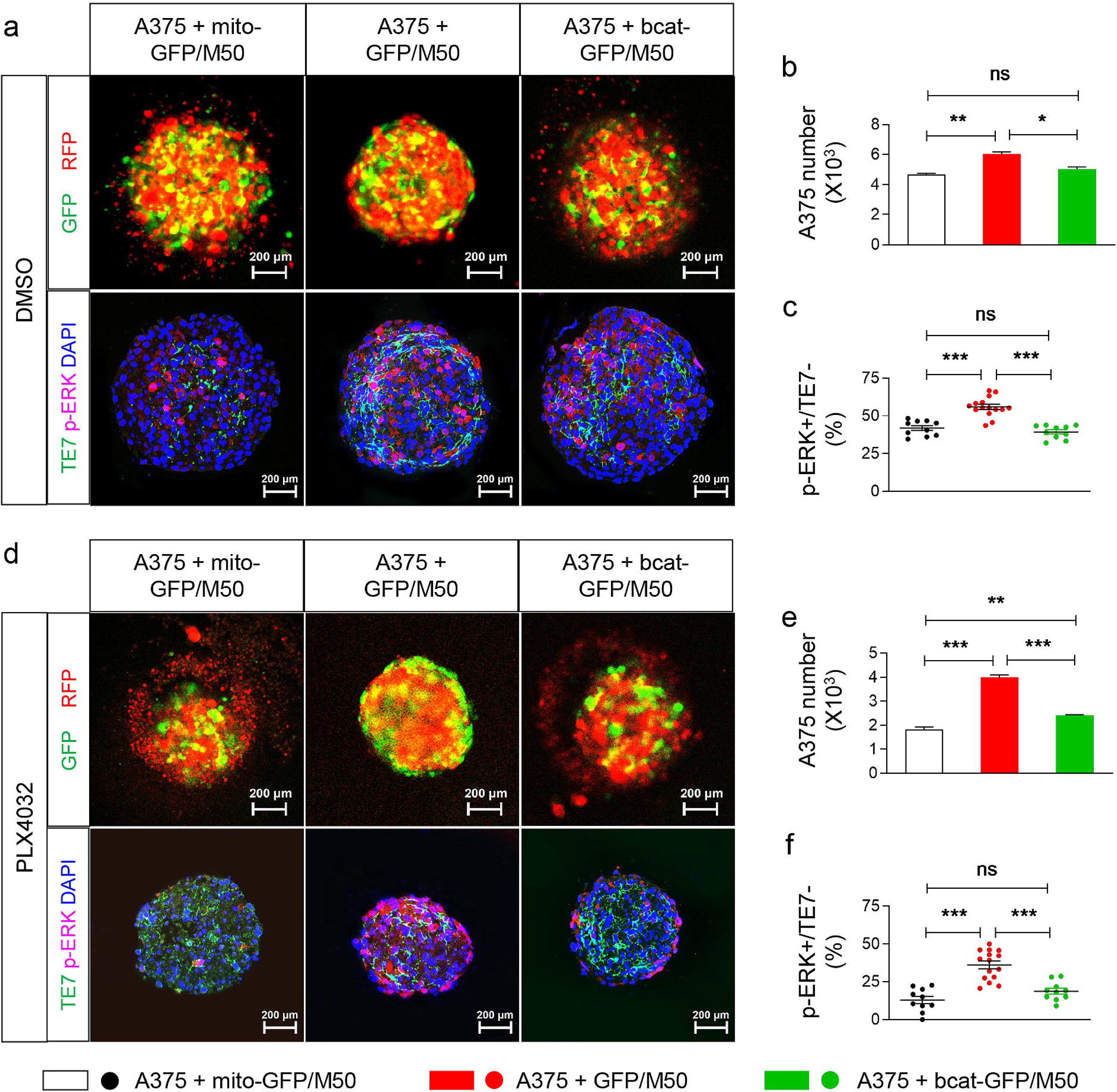
BRAFi-stimulated CAFs require β-catenin to contribute to melanoma cell resistance to BRAFi and MEKi. **a**. Upper panels show fluorescent pictures of 3D spheroid consisting of A375 (red) and M50 (green). Lower panels show immunostaining images using an anti-TE7 antibody (green) and an anti-p-ERK antibody (red) with DAPI counterstaining (blue). Scale bar: 200 μm. **b**. Bar graph shows the mean A375 number per spheroid from three independent experiments. For each group per experiment, 12 spheroids were collected for cell counting, and the number was divided by 12 to obtain the mean A375 number. **c**. Scatter graph shows the percentages of A375 cells (TE7-) expressing p-ERK in each spheroid type. Each data point represents the quantified percentage in each spheroid. n≥10. **d**. Representative images of 3D co-cultured spheroids treated by PLX4032 for 72 hours. Same experimental procedure was performed as described above in DMSO-treated groups. **e**. Bar graph shows the mean A375 numbers per spheroid from three independent experiments. **f**. Scatter graph shows the percentages of A375 cells (TE7-) expressing p-ERK in each spheroid type. Each data point represents the quantified percentage in each spheroid. n≥10. In all graphs, data are represented as mean ± SEM. * *P* ≤ 0.05. ** *P* ≤ 0.01. *** *P* ≤ 0.001. ns: not significant.

When the spheroids were treated with PLX4032 for 72 hours (Fig. 4d, e), there were significantly more melanoma cells in A375 + GFP/M50 spheroids than A375 + mito-GFP/M50 spheroids (119 ± 8 % more A375 cells), showing that M50 cells contributed to the resistance of A375 against PLX4032. β-catenin ablation significantly reduced the ability of M50 cells to elicit the resistance of A375 to PLX4032. The number of A375 in A375 + bcat-GFP/M50 spheroids was only 60 ± 1% of that in A375 + GFP/M50 spheroids. In line with the findings, β-catenin ablation in GFP/M50 reduced the level of p-ERK to 19 ± 2% in A375 when treated with PLX4032 (Fig. 4f). The results were validated using another BRAF^V600E^ melanoma cell line SK-Mel-24 and CAF line M27 (Fig. S3). The data demonstrate that there is a β-catenin dependent crosstalk between melanoma cells and CAFs that allows melanoma cells to bypass the inhibition of BRAFi on MAPK/ERK signaling.

We evaluated whether β-catenin is equally important for CAFs to contribute to melanoma resistance to combination BRAFi/MEKi. As shown in Fig. S4, under PLX4032 and GDC0973 treatment, the number of A375 in the spheroids co-cultured with either M50 or M27 cells were significantly higher than those co-cultured with mitomycin-treated M50 or M27 or β-catenin-deficient M50 and M27.

### Depleting BRAFi-stimulated CAFs sensitizes BRAF-mutant melanoma cells to BRAFi *in vivo*

To investigate whether β-catenin depletion in CAFs has the potential to improve the responsiveness of BRAF-mutant melanoma cells to BRAFi *in vivo*, we established a human melanoma xenograft model using a combination of A375 cells with GFP/M50 or bcat-GFP/M50. As shown in Fig. 5a-c, in severe combined immunodeficiency (SCID) mice fed with saline buffer at day 28, the size and weight of A375 + bcat-GFP/M50 tumor (518.6 ± 36.8 mm^3^; 0.57 ± 0.04 g; n=10) were significantly smaller and lighter than those of control A375 + GFP/M50 tumors (1027 ± 113 mm^3^; 1.03 ± 0.10 g; n=10), confirming that β-catenin activity is important for CAFs to support melanoma progression. Similarly, in the mice administrated with PLX4032 for 28 days, β-catenin-deficient tumors (A375 + bcat-GFP/M50) were also smaller and lighter (222.6 ± 14.2 mm^3^; 0.26 ± 0.02 g; n=11) than those of control A375 + GFP/M50 tumors (549.5 ± 26.6 mm^3^; 0.54 ± 0.02 g; n=11). The results show that ablating β-catenin in CAFs has the potential to increase the responsiveness of BRAF-mutant melanoma cells to BRAFi.

**Fig. 5.**
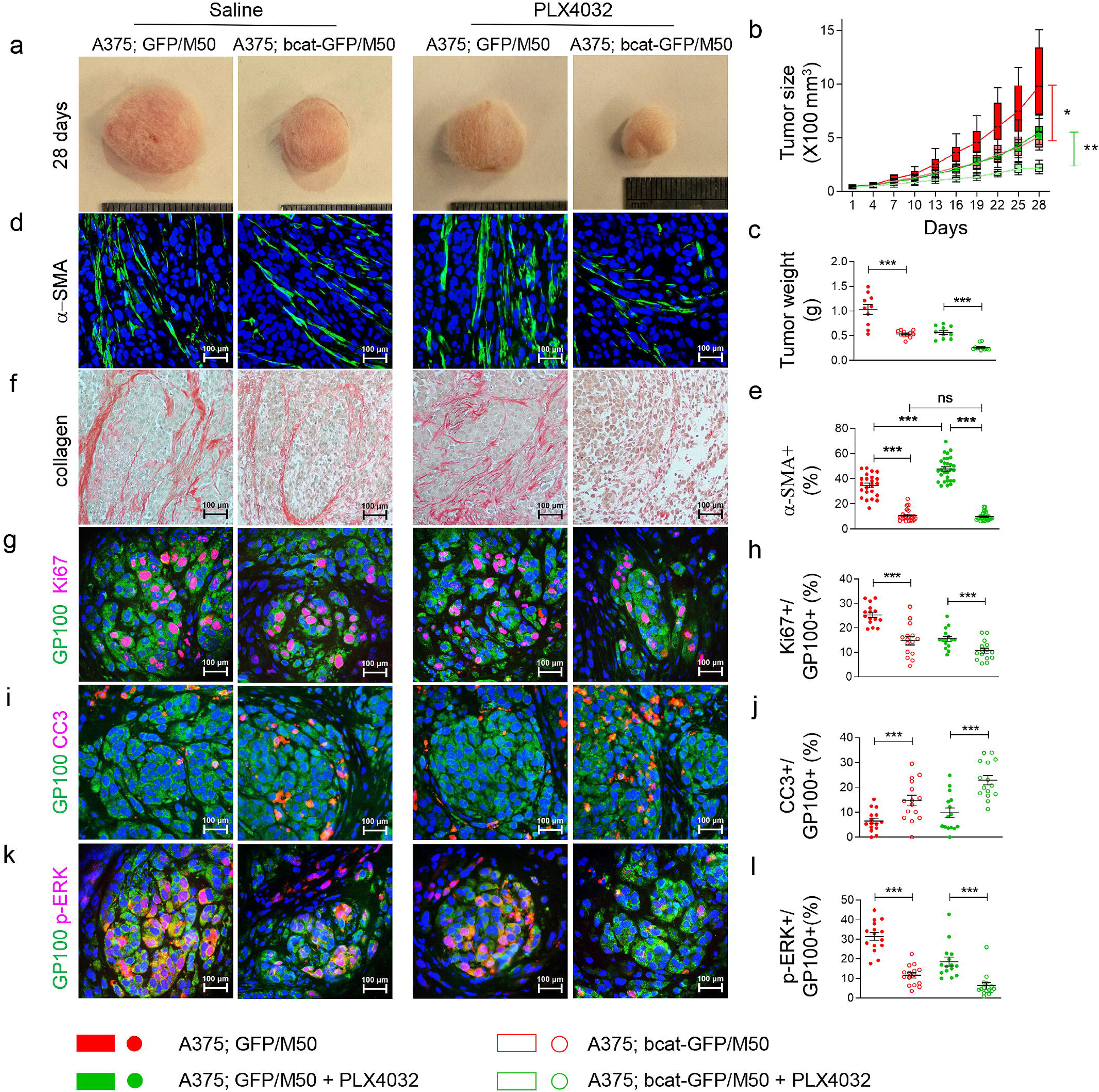
Depleting BRAFi-stimulated CAFs sensitizes BRAF-mutant melanoma cells to BRAFi *in vivo*. **a**. Representative pictures of melanoma xenografts derived from indicated cell mixtures 28 days after saline or Vemurafenib treatment. **b-c**. Comparative analysis of the volumes (**b**) and weights (**c**) of melanoma xenografts between four experimental groups. Box and whisker plot represent the volumes of melanoma xenografts at each indicated time point. Scatter graph shows tumor weight comparison among four groups at day 28. Each data point represents the tumor weight of one tumor. n≥10. **d**. Images show α-SMA staining of melanoma tissues collected from each group. The nuclei were stained blue with DAPI. **e**. Scatter graph shows the percentages of α-SMA+ fibroblasts in total cells from tumors derived from indicated groups. Each data point represents the fibroblast percentage in one 40X field. n≥15. **f**. Images show collagen staining of melanoma tissue sections in indicated groups (red). **g**. Images show GP-100 (melanoma cells, green) and Ki67 (red) staining of melanoma tissue sections in indicated groups. **h**. Scatter graph shows the percentages of GP100+ melanoma cells that are Ki67+ among the groups. Each data point represents the percentage of Ki67+ melanoma cells in one 40X field. n≥15. **i**. Images show GP-100 and cleaved caspase 3+ (CC3+, red) staining of melanoma tissues sections of indicated groups. **j**. Scatter graph shows the percentages of GP100+ melanoma cells that are CC3+ among the groups. Each data point represents the percentage of CC3+ melanoma cells in one 40X field. n≥15. **k**. Images show GP-100 and p-ERK (red) staining of melanoma tissues sections in indicated groups. **l**. Scatter graph shows the percentages of GP100+ melanoma cells that are p-ERK+ among the groups. Each data point represents the percentage of p-ERK+ melanoma cells in one 40X field. n≥15. For all staining pictures: scale bar represents 100 µm. In all graphs, data are represented as mean ± SEM. * P ≤ 0.05; ** P ≤ 0.01; *** P ≤ 0.001. ns: not significant.

As we observed increased fibroblast numbers in post-BRAFi/MEKi treated human melanomas (Fig. 1), we next asked how β-catenin depletion affects the number of M50 in melanoma xenografts. As shown by α-SMA immunostaining in Fig. 5d, e, in the tumors derived from the mixture of A375 + GFP/M50 without PLX4032 treatment, 35 ± 2% of the cells are α-SMA+ fibroblasts. In A375 + bcat-GFP/M50 tumors, the percentage of α-SMA+ fibroblasts was reduced to 11 ± 1%. The reduction is consistent with the reduction in melanoma size and weight. In PLX4032-treated melanomas, the percentage of α-SMA+ fibroblasts in A375 + GFP/M50 tumors was increased to 48 ± 2% and conversely reduced to 10 ± 1% in A375 + bcat-GFP/M50 tumors. CAFs are known to be an important source of fibrillar collagen. PLX4032 treatment promoted collagen production in the melanomas derived from normal CAFs (Fig. 5f). However, a significant reduction of collagen content was observed in the tumors derived from the mixture of A375 + bcat-GFP/M50 with or without PLX4032 treatment.

Next, we investigated the proliferation and apoptosis of melanoma cells (Fig. 5g-l). In A375 + GFP/M50 melanomas without PLX4032 treatment, 25 ± 1% of melanoma cells were Ki67+, and 31 ± 2% of melanoma cells expressed p-ERK. Depleting β-catenin in M50 resulted in a decrease in both the numbers of Ki67+ (15 ± 2%) and p-ERK+ (12 ± 1%) melanoma cells. In A375 tumors carrying GFP/M50 CAFs treated with PLX4032, 16 ± 17% of melanoma cells were Ki67+, and 19 ± 2% of melanoma cells expressed p-ERK. However, in PLX4032 treated A375 tumors containing β-catenin-deficient M50, the percentages of p-ERK+ and Ki67+ cells reduced dramatically to 6 ± 2% and 11 ± 1%, respectively. Cleaved caspase-3 (CC3) is a critical executioner of apoptosis. In mice fed with saline buffer, A375 + GFP/M50 tumors had 7 ± 1% CC3-positive (CC3+) melanoma cells. This percentage increased to 15 ± 2% when β-catenin was depleted in CAFs. In mice treated with PLX4032, the percentages of melanoma cells expressing CC3 increased to 23 ± 2% in A375 + bcat-GFP/M50 tumors, which was significantly higher than the number of CC3+ melanoma cells in A375 + GFP/M50 tumors (10 ± 2%). The data show that β-catenin activity is important for CAFs to contribute to melaoma BRAFi resistance.

As combined BRAF and MEK inhibition has shown promising clinical benefits for individuals with advanced melanoma, we extended the analysis to melanoma xenografts exposed to a combination of BRAFi/MEKi treatment. Smaller tumor size, diminished proliferation, and increased number of apoptotic cells were observed in melanomas carrying β-catenin-deficient CAFs under BRAFi/MEKi treatment (Fig. S5). These findings underscored the importance of BRAFi-induced β-catenin signaling in CAFs in melanoma drug resistance.

### **β**-catenin nuclear translocation is driven by paradoxical activation of RAF kinases in CAFs

**β**-catenin is phosphorylated by glycogen synthase kinase 3 beta (GSK-3**β**) and subsequently ubiquitylated and targeted for proteasomal degradation^32^. As shown in Fig. 6a, in PLX4032-treated M50 cells, the level of phosphorylated GSK-3β (p-GSK-3**β**, marked for GSK-3**β** degradation) displayed a two-fold increase while the amount of p-**β**-catenin (Ser33/37/Thr41, marked for **β**-catenin degradation) decreased significantly. Interestingly, the phosphorylation form of ERK, which is known to have the ability to phosphorylate GSK-3**β**, increased around 6-fold after M50 cells were treated with PLX4032.

**Fig. 6.**
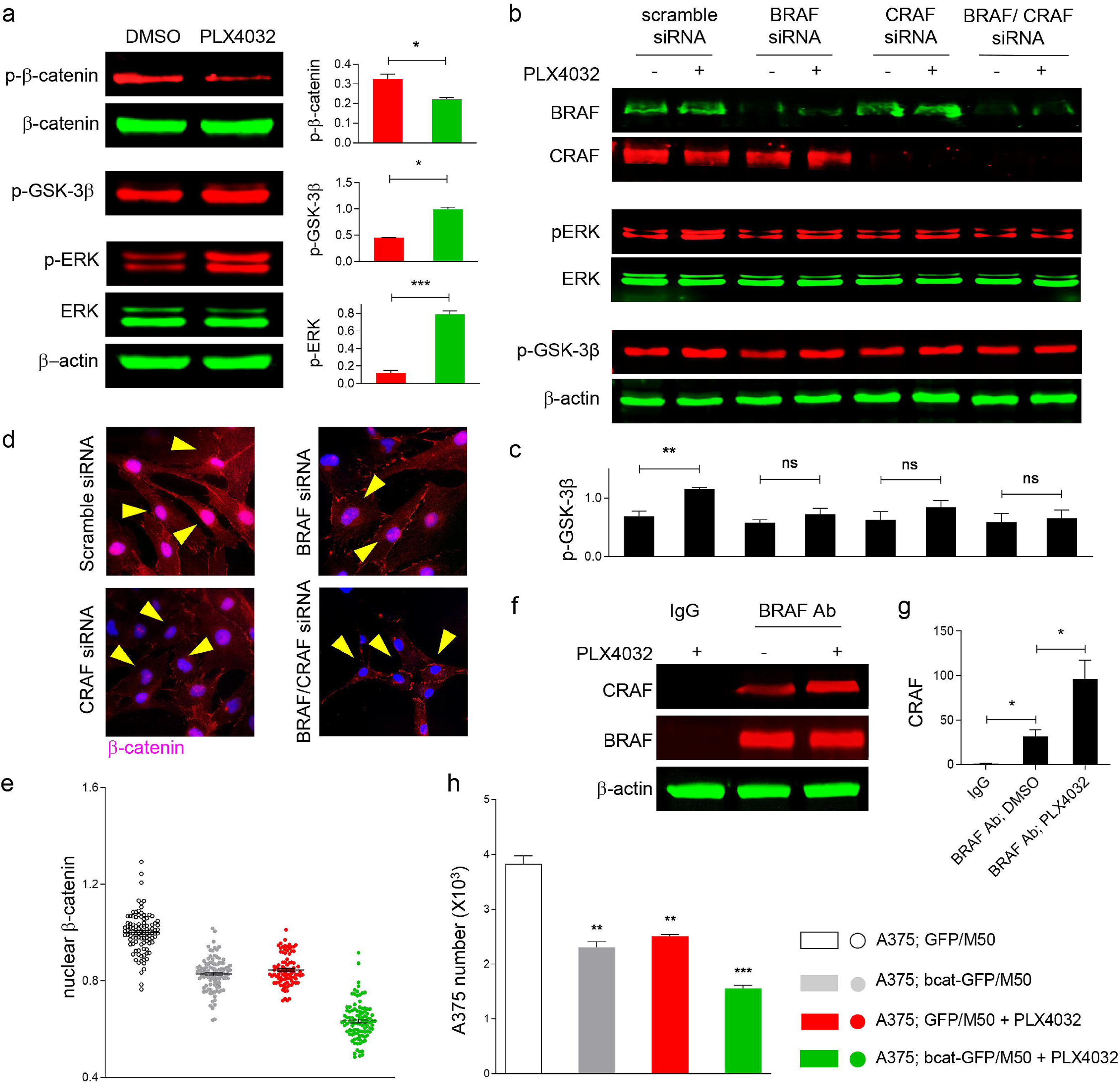
BRAFi-induced BRAF/CRAF dimerization drives β-catenin nuclear accumulation in CAFs. **a**. Western blotting of p-β-catenin (Ser33/37/Thr41), β catenin, p-GSK-3β (Ser9), ERK, and p-ERK levels in M50 treated by DMSO or PLX4032 for two hours. Bar charts show quantification of fluorescence intensities of indicated protein bands after DMSO or PLX4032 treatment. n=3. **b**. Western blotting of BRAF, CRAF, ERK, p-ERK, and p-GSK-3β in M50, in which BRAF, CRAF or a combination of BRAF and CRAF were silenced by siRNAs, treated by DMSO or PLX4032, respectively. **c**. Bar chart shows fluorescent intensities of p-GSK-3β Western blot bands. n=3. **d**. Representative images show β-catenin immunostaining of PLX4032-treated M50 with silenced BRAF or CRAF or BRAF/CRAF expression (indicated by yellow triangles). **e**. Scatter graph shows relative nuclear β-catenin expressions in M50 treated with PLX4032 after BRAF, CRAF or a combination of BRAF and CRAF were silenced by siRNAs. The expression level of nuclear β-catenin was normalized by the mean nuclear β-catenin expression in M50 transfected by scramble siRNA after a 72-hour PLX4032 treatment. n≥ 0. **f**. Western blotting of BRAF and CRAF in co-immunoprecipitated M50 samples treated by DMSO or PLX4032 for 72 hours using IgG or an anti-BRAF antibody. **g**. Bar chart shows fluorescence intensities of CRAF bands from indicated co-IP samples. n=3. **h**. Single or double knock down of BRAF and CRAF partially deprives of the ability of CAFs to promote A375 resistance to PLX4032. Bar chart shows the numbers of A375 in the spheroids co-cultured with M50 with silenced BRAF or CRAF or BRAF/CRAF expression after a 72-hour PLX4032 treatment. n=3. For all graphs: data are shown as mean ± SEM. * *P* ≤ 0.05. ** *P* ≤ 0.01; *** *P* ≤ 0.001. ns: not significant.

It has been shown that ATP-competitive RAF inhibitors can bind to and activate wildtype BRAF and consequently activate the downstream signaling cascades^33–37^. Therefore, we hypothesized that increased nuclear β-catenin in BRAF-wildtype CAFs is driven by BRAFi-induced BRAF and CRAF dimerization and activation of the downstream ERK signaling pathway. The expression of BRAF, CRAF, or a combination of BRAF and CRAF was silenced in CAFs by siRNAs. Immunoblotting results in Fig. 6b show that the expression of BRAF or/and CRAF was successfully knocked down in M50 cells. Ablation of either BRAF or CRAF in M50 was able to reduce PLX4032-induced increases in p-ERK and p-GSK-3β levels, indicating that both BRAF and CRAF activities are required for PLX4032-induced nuclear β-catenin accumulation in M50 cells. Depleting both BRAF and CRAF blocked p-ERK and p-GSK-3β increase in M50 (Fig. 6c). The important roles of BRAF and CRAF were further underscored by the fact that either BRAF or CRAF silencing greatly suppressed BRAFi-induced β-catenin nuclear translocation, and ablation of both BRAF and CRAF was sufficient to diminish nuclear β-catenin accumulation in

M50 induced by PLX4032 (Fig. 6d, e). Co-immunoprecipitation was performed to determine whether BRAFi leads to BRAF/CRAF dimerization in CAFs. An anti-BRAF antibody was used to bait CRAF protein from the cell lysate of M50 cells with or without PLX4032 treatment. The binding of BRAF and CRAF could be observed in M50 cells in the absence of any inhibitor (Fig. 6f). PLX4032 treatment led to a two-fold increase of BRAF/CRAF dimer formation (Fig. 6g). The data demonstrate that PLX4032 promotes the formation of a heterodimer between BRAF and CRAF in CAFs and leads to increased ERK signaling and GSK-3β phosphorylation. 3D co-culture assays also showed that both BRAF and CRAF are required by CAFs to contribute to melanoma cell resistance to PLX4032 (Fig. 6h). Altogether, the above findings establish that BRAFi drives the activation of RAF kinases and elicits nuclear β-catenin signaling in CAFs.

To confirm that ERK signaling is a major pathway downstream RAF activation that contributes to β-catenin nuclear accumulation in CAFs, we used ERK1/2 inhibitor SCH772984 to treat PLX032-treated M50^38^. Surprisingly, PLX4032-induced β-catenin nuclear accumulation in M50 was blocked by SCH772984 (Fig. S6a, b). Furthermore, the increase in the level of p-GSK-3β induced by PLX4032 was also inhibited by SCH772984 (Fig. S6c, d). Altogether, the above findings establish that BRAFi drives the activation of nuclear β-catenin signaling via RAF kinases and ERK signaling in CAFs.

### POSTN is a downstream effector of **β**-catenin signaling in CAFs

To obtain a global picture on the underlying mechanisms by which β-catenin regulates the functional properties of CAFs and identify β-catenin-regulated genes that are involved in CAF-elicited melanoma BRAFi/MEKi resistance, we isolated wildtype CAFs and β-catenin-deficient CAFs (bcat/CAFs) from the BRAF-mutant melanoma-CAF mouse model that were developed in our laboratory^25^ for RNA-Seq (NCBI GEO Accession number: GSE121186, review token gnsbemawnzizxaz). Normal fibroblasts (NFs) isolated from C57BL/6J mouse skin were used as a control. The genes that were expressed in CAFs two-fold above or below the corresponding ones in both NFs and bcat/CAFs were identified. This strategy yielded a filtered list of 1790 genes as plotted in the heatmap (Fig. 7a). Significant changes in gene expression were observed in wildtype CAFs compared with normal fibroblasts; however, β-catenin ablation in CAFs dramatically altered the gene expression pattern, which resembled the one of normal fibroblasts. The data suggest that β-catenin is a major driver of the biological identity of CAFs, and depleting β-catenin reverts CAFs into a normal-like fibroblast state. Using Venn diagrams39, we compared genes that were upregulated or downregulated at least five times in either NF or bcat/CAFs in comparison to CAFs. In the total 1054 genes filtered, a 79.9% overlap between significantly changed genes in CAFs (NFs as a control) and bcat/CAFs (CAFs as a control) were observed (Fig. 7b), further confirming that bcat-deficient CAFs phenocopied NFs. The genes that were reduced at least two-fold in bcat/CAFs compared with CAFs underwent a functional annotation clustering analysis using DAVID Bioinformatics Resources 6.8 (Fig. 7c)^40^. Most of those differentially expressed genes fall into the categories of ECM production, focal adhesion, paracrine signaling, intracellular signaling, transcription regulation, and cytoskeleton remodeling.

**Fig. 7.**
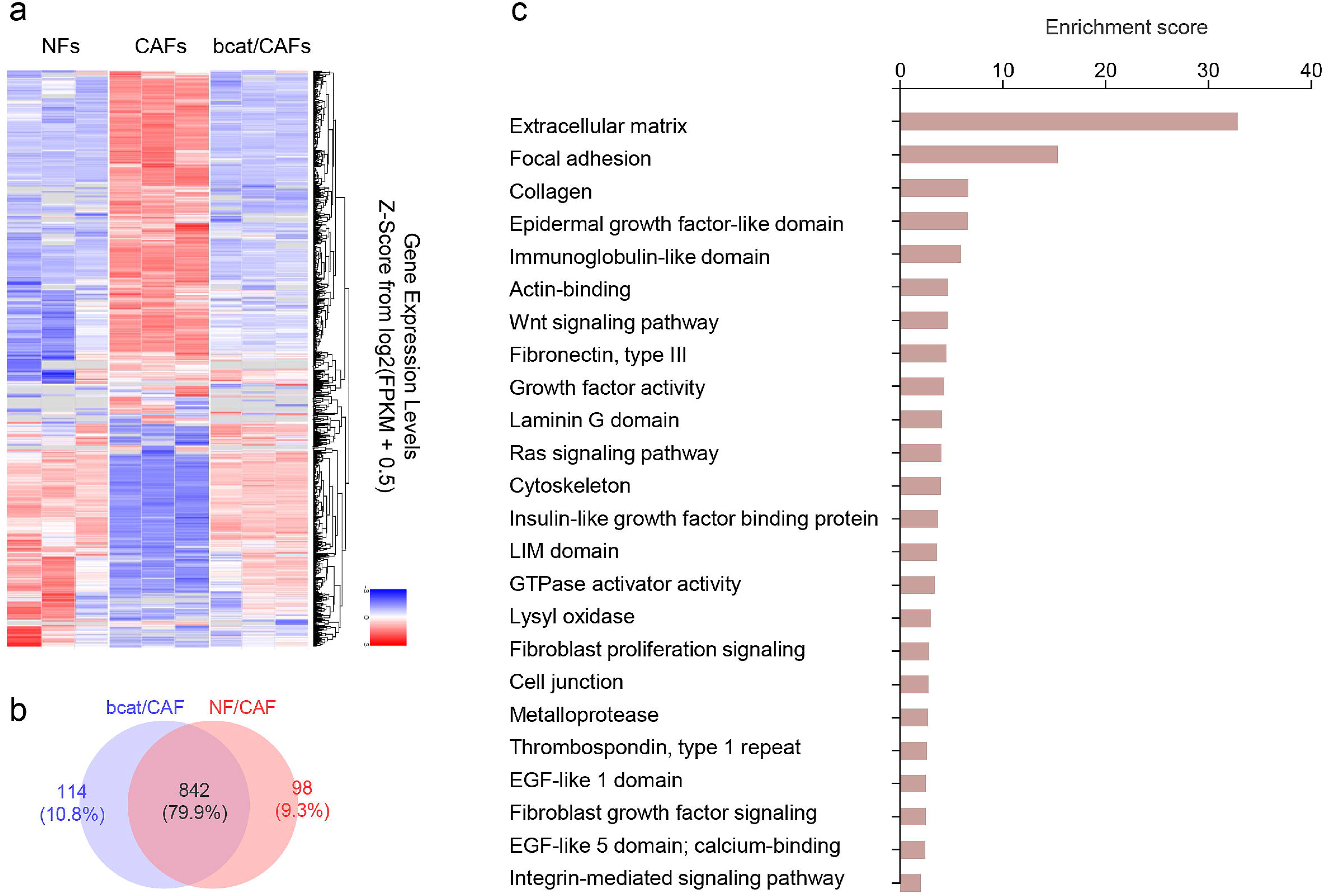
β-catenin is a master regulator in CAFs. **a**. Heatmap shows the top 1790 most upregulated and downregulated genes among NFs, CAFs, and bcat/CAFs. **b**. Venn diagram compares genes that were upregulated or downregulated at least five times in either NF or bcat/CAFs in comparison to CAFs. **c**. The graph shows the functional annotation clustering analysis of differentially expressed genes that are down-regulated more than two times in bcat/CAFs using DAVID Bioinformatics Resources 6.8. Enrichment Score for each group indicates the biological significance of gene groups based on overall EASE scores of all enriched annotation terms.

Based on this list of genes, we identified POSTN as a top downstream effector of β-catenin that mediates CAF-induced melanoma drug resistance. To evaluate the significance of POSTN for BRAFi resistance, we performed paired analyses of POSTN expression in melanoma tissues isolated from the same melanoma patients before and after BRAFi/MEKi therapy (Fig. 8a, b) and found that the majority of POSTN-expressing cells were also positive for TE7 expression^31^, suggesting that POSTN was mainly produced by fibroblasts in melanoma. Furthermore, the percentage of POSTN-expressing fibroblast in TE7+ fibroblasts increased from 36 ± 2% (pre-treatment) to 47 ± 2% (post-treatment). *In vitro* validation experiments confirmed that POSTN is highly expressed in all CAF lines we have procured but not in any melanoma cell lines we tested, including A375, SK-MEL-24, 1205Lu, and WM278 (Fig. 8c). In addition, the POSTN expression was upregulated by PLX4032 in CAFs, while β-catenin depletion blocked PLX4032-induced POSTN upregulation (Fig. 8d).

**Fig. 8.**
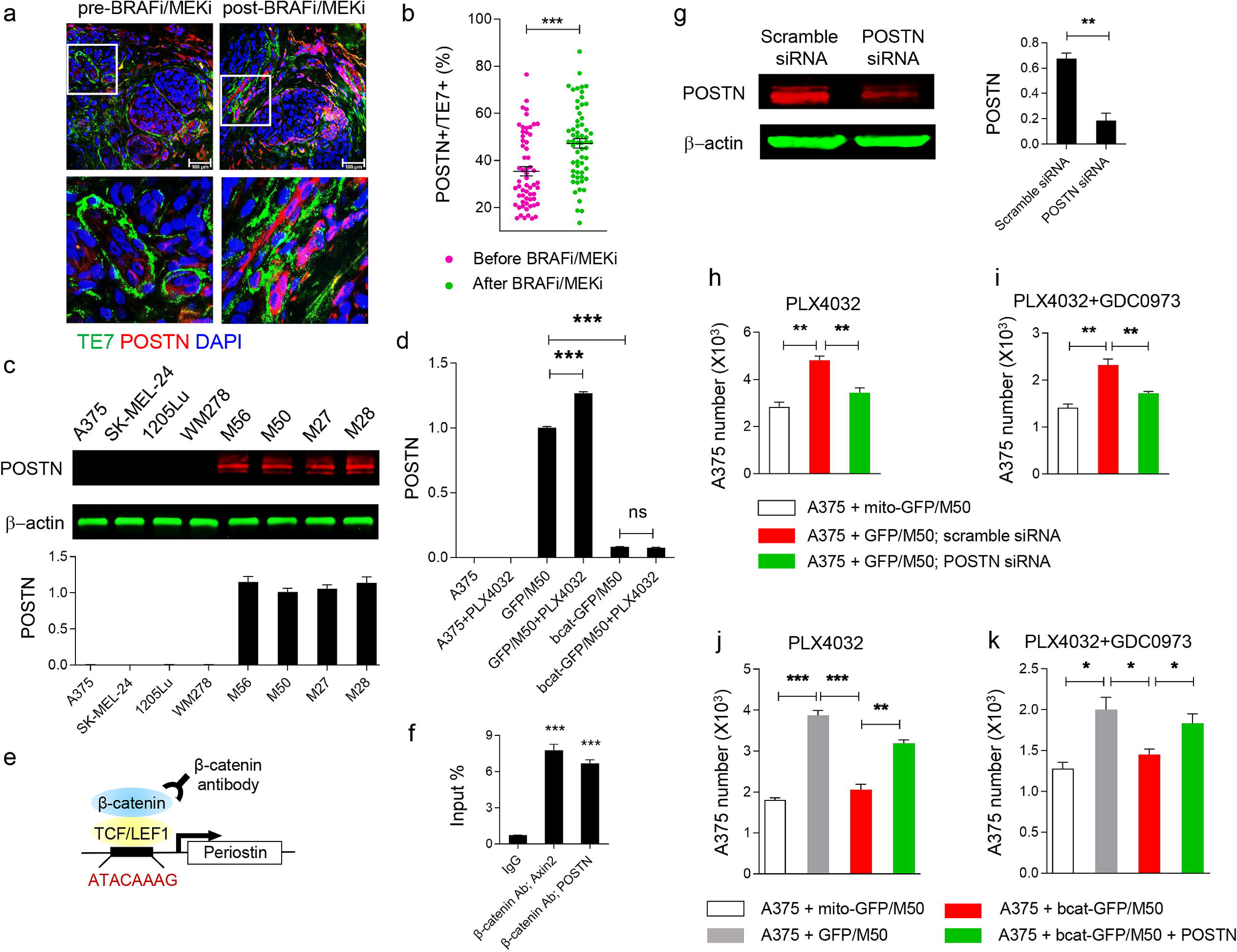
POSTN is a direct downstream effector of nuclear β-catenin signaling in CAFs. **a**. Fibroblasts and POSTN expression in human melanomas pre- and post-BRAFi/MEKi treatment were visualized by co-immunostaining using an anti-TE7 antibody (green) and an anti-POSTN antibody (red). Scale bar: 100 μm. **b**. Scattered plot represents the percentage of TE7+ cells expressing POSTN in pre- and post-treated melanoma samples. Each data point represents the percentage of TE7+ fibroblasts that are POSTN+ in each 40X field. n≥ 40. **c**. Western blotting of POSTN expression levels in indicated melanoma cells and CAFs. Bar chart shows fluorescence intensities of POSTN bands. n=3. **d**. Bar graph shows qPCR data of POSTN expression in A375, GFP/M50, and bcat-GFP/M50 treated with DMSO or PLX4032. n=4. **e**. Illustration of the promoter region of periostin that contains a potential LEF/TCF-binding site. **f**. Bar graph shows signals obtained from ChIP normalized by signals obtained from the input sample. Axin 2 was used as a positive control. n=3. **g**. Western blotting of POSTN expression in M50 transfected with scramble siRNA or POSTN siRNA. Bar graph shows fluorescence intensities of POSTN bands from three independent experiments. **h-i**. Bar graph shows A375 numbers in 3D co-cultured spheroids of indicated groups under PLX4032 treatment (**h**) or PLX4032 and GDC0973 treatment (**i**). n=3. **j-k**. Bar graph shows A375 numbers in 3D co-cultured spheroids of indicated groups with or without recombinant POSTN under PLX4032 treatment (**j**) or PLX4032 and GDC0973 treatment (**k**). n=3. For all graphs, data are shown as mean ± SEM. * *P* ≤ 0.05. ** *P* ≤ 0.01; *** *P* ≤ 0.001. ns: not significant.

### POSTN is a direct β-catenin target in CAFs

We identified an evolutionarily conserved TCF/LEF-binding site in the proximal promoter region of the POSTN gene (chr13:37599016-37599023, 2009 human reference sequence GRCh37, Fig. S7) using PROMO^41, 42^ (Supp file 1). Guided by this observation, a chromatin immunoprecipitation (ChIP) assay was performed to investigate whether POSTN is a direct β-catenin target in CAFs (Fig. 8e). As shown in Fig. 8f, significant binding of β-catenin was found in the POSTN promoter region that contains the TCF/LEF consensus sequence. We then used siRNA to silence POSTN expression in M50 cells (Fig. 8g). 3D drug resistance assay using PLX4032 or combined PLX4032 + GDC0973 showed that fewer melanoma cells survived therapy treatment in the spheroids co-cultured with POSTN-deficient M50 cells compared with control M50 cells (Fig. 8h, i), indicating that POSTN is required for CAF-mediated drug resistance to BRAFi/MEKi.

To determine whether POSTN can compensate for the loss of β-catenin in CAFs and induce melanoma resistance to BRAFi/MEKi, we added recombinant POSTN to the spheroids formed by A375 and bcat-GFP/M50 under single PLX4032 or combined PLX4032 + GDC0973 treatment (Fig. 8j, k). The data show that recombinant POSTN, at least partially, rescues the loss of β-catenin in CAFs and promotes melanoma resistance to BRAFi and MEKi.

### POSTN reactivates ERK signaling in BRAF-mutant melanoma cells to bypass BRAFi and MEKi inhibition

A375 cells grew better and had increased resistance to PLX4032 when cultured on the plates coated with recombinant POSTN (Fig. 9a-c). We confirmed these findings using 3D melanoma spheroids (Fig. 9d-f). Recombinant POSTN induced an increase in melanoma spheroid size. Furthermore, recombinant POSTN also induced A375 resistance to MEKi. As shown in Fig. 9g, A375 gained resistance to GDC0973 and the combination of PLX4032 and GDC0973 when POSTN was added.

**Fig. 9.**
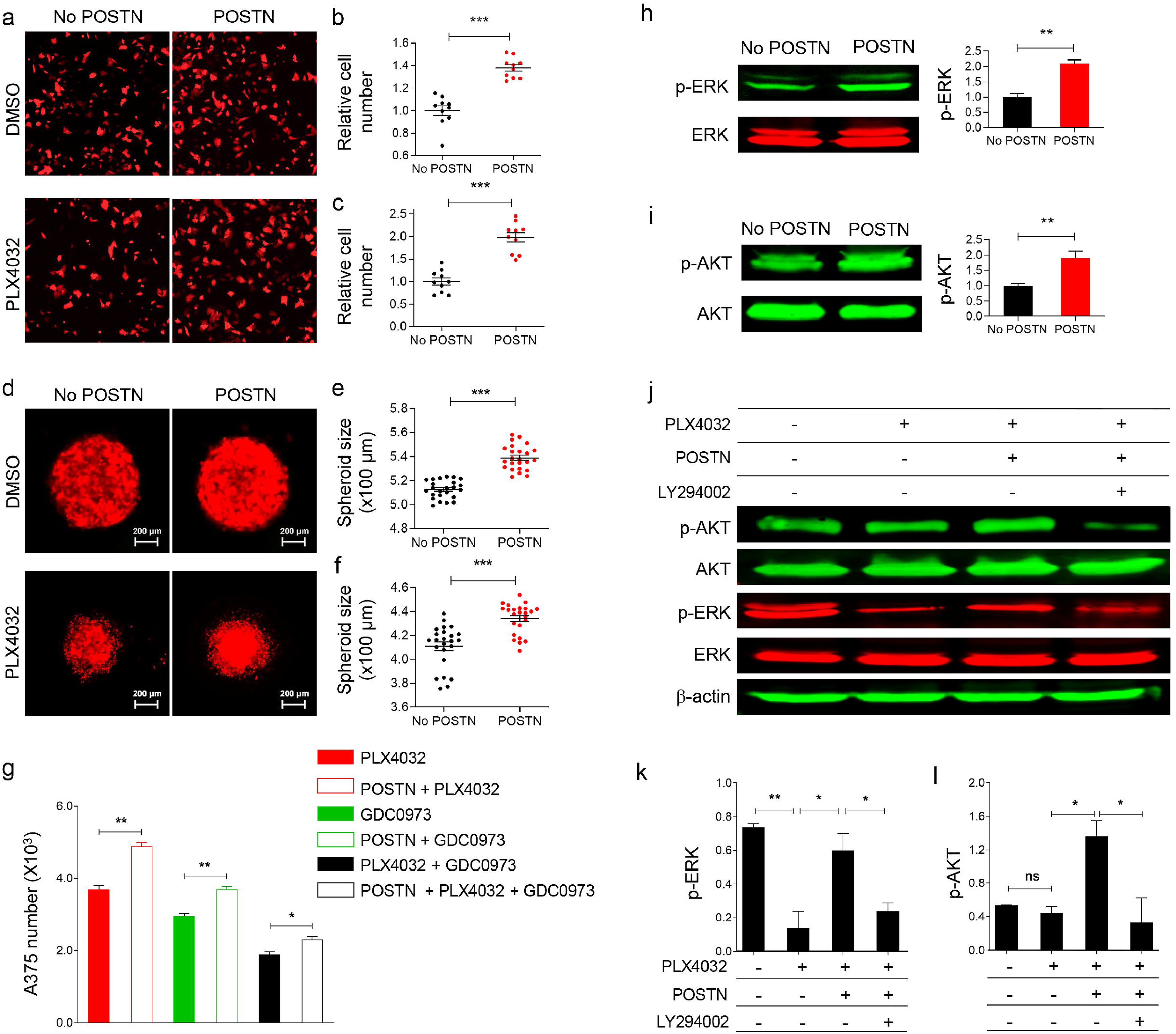
POSTN activates alternative PI3K/AKT signaling in BRAF-mutant melanoma cells to bypass BRAFi and MEKi inhibition. **a**. Fluorescent images show RFP-tagged A375 cells in collagen-coated dishes and collagen + POSTN-coated dishes with or without PLX4032 treatment. **b-c**. Scatter graph shows relative A375 numbers in regular and POSTN-coated dishes under DMSO (**b**) or PLX4032 (**c**) treatment. Each data point represents relative cell number in each 4X field normalized by the mean A375 number in No POSTN group. n=10. **d**. Representative images of 3D RFP-tagged A375 spheroids cultured with and without POSTN under DMSO or PLX4032 treatment. Scale bar: 200 µm. **e-f**. Scatter graph shows the sizes of the spheroids cultured with or without POSTN under DMSO (**e**) or PLX4032 treatment (**f**). Each data point represents the size of one spheroid from indicated groups. n=24. **g**. Bar graph shows the numbers of A375 cultured with and without POSTN when treated with PLX4032, GDC0973, or a combination of PLX4032 and GDC0973. n=3. **h**. Western blotting of ERK and p-ERK levels in A375 treated with and without POSTN. Bar chart shows fluorescent intensities of ERK and p-ERK bands. n=3. **i**. Western blotting of AKT and p-AKT levels in A375 treated with and without POSTN. Bar chart shows fluorescent intensities of AKT and p-AKT bands. n=3. **j**. Wester blotting of AKT, p-AKT, ERK, and p-ERK levels in A375 treated with PLX4032, POSTN, and LY294002 as indicated in the figure. **k-l**. Bar chart shows fluorescence intensities of p-ERK bands (**k**) and p-AKT bands (**l**) under indicated conditions. n=3. For all graphs, data are represented as mean ± SEM. * *P* ≤ 0.05. ** *P* ≤ 0.01; *** *P* ≤ 0.001. ns: not significant.

POSTN is an ECM protein that establishes contact and communication between the ECM microenvironment and the cell via binding to integrin receptors^43^. It was reported that POSTN activates the integrin/AKT/MAPK pathway in cancer cells to accelerate cell proliferation^44^. Given that the addition of POSTN promoted melanoma growth and drug resistance, we collected POSTN-treated A375 and investigated the phosphatidylinositol 3 kinase (PI3K)/AKT pathway and ERK pathway. Interestingly, a roughly two-fold increase was observed in the expression of p-ERK and p-AKT in POSTN-treated A375, suggesting that AKT and ERK signaling were activated (Fig. 9h, i). To confirm whether POSTN induces the reactivation of ERK signaling in BRAFi-treated A375 and whether this alternative pathway is PI3K/AKT-dependent, we assessed the activation of the AKT and ERK pathways in A375 treated with POSTN and PLX4032 (Fig. 9j-l). In A375 treated with PLX4032 only, p-ERK was significantly inhibited while the p-AKT level remained unchanged compared with control A375 cells. However, the addition of POSTN not only restored the level of p-ERK that was inhibited by PLX4032 but also led to a significant increase of p-AKT level in PLX4032-treated A375 cells. LY294002 is known as a highly selective PI3K inhibitor^45^ and was added to inhibit AKT phosphorylation by PI3K in A375. POSTN-induced reactivation of ERK signaling in A375 under PLX4032 treatment was blocked by LY294002. The data show that POSTN reactivates the ERK signaling pathway in melanoma cells, which is inhibited by BRAFi and MEKi, through PI3K/AKT signaling.

## DISCUSSION

Melanomas are solid tumors that harbor a significant number of stromal cell populations. The myriads of interactions between melanoma cells and the stromal components mold many core characteristics of cancer, the so-called hallmarks^7^. However, for the longest time, the main thrust in cancer therapy has been to exploit “weaknesses” of cancer cells themselves, such as oncogene addiction, high proliferation rates, or specific mutations in suppressor genes. Such efforts have resulted in the development of BRAFi that specifically targets the frequent BRAF mutations in melanoma cells. Nevertheless, the development of drug resistance after the initial positive response is the biggest hurdle to achieve the ultimate success of BRAF-targeted cancer therapies. The data reported here highlight that the occurrence of drug resistance is not a simple response by merely melanoma cells and can substantially be affected by complex interactions among tumor cells, stromal cells, the ECM, and targeted therapy drugs (Fig. 10).

**Fig. 10.**
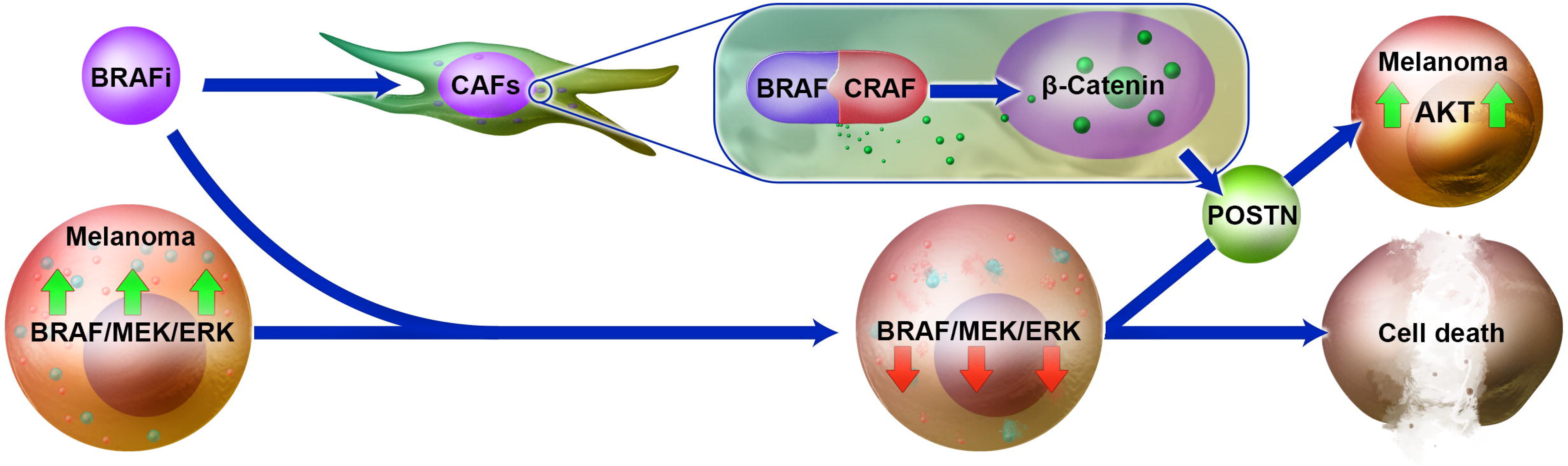
Model of BRAFi-induced CAF reprogramming in matrix remodeling and melanoma BRAFi/MEKi resistance. BRAFi blocks BRAF/MEK/ERK signaling in BRAF V600E mutant melanoma cells, leading to cell death. On the other hand, BRAFi stimulates CAFs and elicits the formation of BRAF/CRAF dimers and nuclear β-catenin accumulation. Nuclear β-catenin signaling reprograms CAFs and leads to the production of matricellular protein POSTN. POSTN activates alterative PI3K/AKT signaling in BRAF-mutant melanoma cells and subsequently ERK activation, which allows melanoma cells to bypass BRAFi/MEKi.

Our study demonstrates that CAFs play important roles in supporting melanoma cell survival and proliferation under targeted therapy. CAFs are not a static population of fibroblasts and possess significant heterogeneity in characteristics and functions, which are in fact a direct reflection of their interactions with the surrounding environment^20^. Several studies reported that distinct CAF populations exist within a single tumor based on their gene expression profiles^22, 46^. This is clearly demonstrated by the heterogeneity of six CAF cell lines used in our study, displaying different levels of α-SMA, F-actin, paxillin, and p-MLC2 expression. In addition, these CAF cell lines show strong but variable abilities to migrate and contract collagen gels. Importantly, we discovered that BRAFi led to an expanded α-SMA+ CAF population with increased nuclear β-catenin accumulation in melanoma. These α-SMA+/nuclear β-catenin+ CAFs have enhanced biological properties, especially their abilities to remodel the cytoskeleton and the ECM. The data suggest that CAFs in BRAFi/MEKi-treated melanoma develop the characteristics of hyperactivated ECM-remodeling myofibroblast-like CAFs. Furthermore, the findings reveal that BRAFi is able to elicit adaptive transcriptional and cellular activities in CAFs and equip them with specific phenotypes that allow them to survive better under the pressure of therapeutic agents and help melanoma cells to escape BRAFi.

Conventionally, nuclear β-catenin accumulation is associated with canonical Wnt signaling^47^. Under the normal circumstance, without Wnt ligands, cytoplasmic β-catenin is phosphorylated at its N-terminus by GSK-3β in a destruction complex containing the adenomatous polyposis coli protein (APC) and Axin for degradation via the ubiquitin/proteasome pathway. When the Wnt pathway is activated by Wnt ligands, GSK-3β is phosphorylated, and β-catenin is stabilized and translocated into the nucleus. However, in our study, we have not found any evidence showing the increased production of Wnt ligands by either melanoma cells or stromal cell populations. Instead, our data suggest that CAFs also succumb to the “paradox” effects of BRAFi, showing BRAF and CRAF dimerization, which results in the activation of ERK signaling and nuclear β-catenin accumulation. Wnt-independent activation of nuclear β-catenin signaling has been reported before^48^, e.g., drugs that target the proteasome, thereby stabilizing β-catenin protein^49^. In 2001, *Morali et al* reported that stimulation of epithelial cells with IGF2 could induce nuclear β-catenin accumulation mainly through the degradation of E-cadherin (CDH1) and re-distribution of β-catenin to the nucleus^50^. Another similar mechanism underlying Wnt-independent β-catenin activation involves cadherin internalization, leading to the release of β-catenin from the membrane pool^51, 52^. However, in our case, it appears that BRAFi-induced nuclear β-catenin activation in CAFs is likely independent of cadherins, especially siliencing BRAF and CRAF in CAFs effectively inhibited the BRAFi-induced nuclear β-catenin accumulation.

BRAFi, such as PLX4032, selectively interferes with the kinase domain of BRAF molecules carrying a V600E mutation and inhibits its activity^53^. Therefore, BRAFi is enormously effective in controlling melanomas that have not developed widespread resistance towards the drugs. However, such ATP-competitive BRAFi has paradoxical side effects on the activation of wildtype RAF signaling^54, 55^. It was reported that BRAFi has the ability to induce kinase domain dimerization among the wildtype members of the RAF family, for example, BRAF/CRAF dimers^36, 37^, and such dimerization is a critical event in RAF activation^56^. So far, most of the observed “paradoxical” side effects of BRAFi have been reported to occur in primary cutaneous malignant lesions^57, 58^. The fact that we observe an increase in CRAF/BRAF dimerization in CAFs in response to BRAFi provides evidence that BRAFi alters not just BARF-mutant melanoma cell behavior but the behavior of stromal cell types carrying wildtype BRAF. Eventually, the BRAFi-mediated BRAF/CRAF dimerization results in the activation of MEK-ERK signaling and the nuclear localization of β-catenin in CAFs carrying wildtype BARF.

The finding in this report could reasonably explain why a CAF population with increased nuclear β-catenin is expanded in melanoma after BRAFi treatment. Nuclear β-catenin binds the members of the T-cell factor/lymphoid enhancer factor (TCF/Lef1) family and activates its target gene expression^59^. Resulting β-catenin transcriptional activities have been demonstrated in fibroblast activation, fibrosis, and tissue repair^60–62^. As such, BRAFi-induced β-catenin nuclear activation in CAFs has a plethora of consequences beyond the impact of the canonical function of RAS-RAF-MEK-ERK activation and drives a program in CAFs that enhances (1) their biological functions; (2) their ability to contract collagen and stiffen the ECM and (3) their ability to protect melanoma cells from the toxic effects of BRAFi. Our transcriptome analysis supports this finding and shows that a loss of β-catenin completely reverses the CAF transcriptome into a transcriptome that resemble normal fibroblasts. This astonishing transcriptomic reversal of CAFs into normal fibroblasts reveals a previously unappreciated but pivotal role of β-catenin for CAF functions. The changes are of such an extent that β catenin can be regarded as a master regulator of the CAF phenotype that is modulated by BRAFi. The fact may also explain why the CAF subset with increased nuclear β-catenin is expanded in BRAFi-treated melanoma. In addition, there is a possibility that β-catenin could eventually be considered to be a biomarker for BRAFi-stimulated CAFs although more clinical validation and tests are required.

Our analysis has focused on a top β-catenin target gene in CAFs, POSTN, which is known as an important matricellular protein that is linked to human malignant melanoma^44, 63–65^. Our study illustrates the importance of POSTN as it is only highly expressed in CAFs in melanoma tissues, and the production of POSTN is increased in CAFs in BRAFi-treated melanoma tissues. In addition, PLX4032 increases POSTN production in CAFs but fails to show the same effect in β-catenin-deficient CAFs. Furthermore, we confirm that POSTN is a β-catenin target in CAFs by the ChIP assay, and β-catenin-deficient CAFs do not produce POSTN. Most importantly, POSTN partially rescues β-catenin-deficient CAFs in promoting melanoma cell proliferation and BRAFi/MEKi resistance. POSTN may have multiple effects on melanoma cells, directly interacting with integrin receptors on the cell membrane or indirectly remodeling the ECM structure. In our study, we discovered that POSTN directly induces an alternative PI3K/Akt pathway in melanoma cells and leads to the activation of ERK signaling, allowing melanoma cells to continue to proliferate despite the dual inhibition by BRAFi and MEKi on RAF/MEK signaling (Fig. 10). The data indicate that POSTN could be an ideal target to disrupt the tumor microenvironment and improve targeted therapy or even immunotherapy. In fact, multiple angles have been explored to suppress POSTN^66^, including targeting POSTN mRNA with siRNAs^67–69^, or targeting POSTN protein with anti-POSTN antibodies^70, 71^ or DNA aptamers^72^ although a combination of POSTN targeting and targeted therapy or immunotherapy has not been tried and may worth further investigation.

Our study presents novel insights into the biology of CAFs, their adaptive response to targeted therapy drugs, and how CAFs and CAF-derived signals may be targeted to improve current therapies. We demonstrated that β-catenin is a master regulator of the BRAFi-stimulated CAF phenotype. Identifcaiton of its downstream effectors, such as POSTN, offers a novel angle to attack an important cancer-supporting cell type and break the microenvironment, in which tumor cells reside. Taken together, the data reveal a signaling loop among BRAFi, CAFs, the matrix, and tumor cells that drives the development of acquired resistance in melanoma.

## MATERIALS AND METHODS

### Human melanoma samples

Human melanoma tissues sections were provided by the University of Cincinnati Biorepository and the Center for Rare Melanomas at the University of Colorado. All melanoma tissue samples were previously obtained by both institutions and were not collected specifically for this study.

### Cell lines

Human melanoma cell lines, A375 [*BRAF(V600E)*; *PTEN(mut)*] and SK-MEL-24 [*BRAF(V600E); PTEN(mut)*; *CDKN2A*], were purchased from American Type Culture Collection (ATCC, Manassas, VA) and maintained in Dulbecco’s modified Eagle medium (DMEM) in a humidified incubator at 37°C with 5% CO_2_. Four CAF lines (DT01056P1, 224350P1, DT01027P1, DT01028P1) isolated from surgically excised human melanoma tissues were purchased from Asterand bioscience (Detroit, MI) and designated as M56, M50, M27, M28, respectively. CAF M77 and M80 were isolated and provided by Dr. Guangyong Peng at the Saint Louis University School of Medicine. Human dermal fibroblasts were provided by Dr. Ana Luisa Kadekaro at the Department of Dermatology of the University of Cincinnati. Human fibroblasts and CAFs were maintained in DMEM in a humidified incubator at 37°C with 5% CO_2_. All culture media were supplemented with 10% (v/v) fetal bovine serum (FBS), 1% (v/v) 10,000 U/ml penicillin, and 10,000 U/ml streptomycin. All cell culture reagents were purchased from Thermo Fisher Scientific (Rochester, NY) unless otherwise stated. All experimental procedures involving biosafety issues were carried out under the University of Cincinnati Institutional Biosafety Committee protocol 16-08-17-01.

### Lentiviral shRNA

To stably silence β-catenin expression, we transduced CAFs, including M27 and M50, with inducible lentivirus expressing shRNA that specifically targets β catenin expression (GE Dharmacon, Lafayette, CO, cat# V3SH7675-02EG1499^24^). Inducible lentivirus expressing non-target shRNA (GE Dharmacon, cat# VSC6572) was used to generate control CAFs. The lentiviral vector carries genes encoding a puromycin-resistance protein and GFP. Thus, transduced CAFs can be selected by puromycin. The inducible lentiviral shRNA vector, which utilizes the Tet-On inducible system, only allows the expression of either β-catenin-targeting shRNA or non-target shRNA when cells are treated with doxycycline. Upon induction, GFP expression is simultaneously induced in CAFs so that the cells expressing shRNA can be visually tracked by green fluorescence. To distinguish melanoma cells from CAFs, non-target shRNA RFP (GE Dharmacon, cat# VSC6573) was used to tag melanoma cells with red fluorescence.

Briefly, the cells were first seeded in 6-well tissue culture plates to reach 50% confluence, and then 10 µl of 1 x 10^7^ TU/ml lentiviral particles and 5 µg/ml polybrene (American Bioanalytical, Cat #AB01643) in 500 µl of Opti-MEM I Reduced Serum Medium (Thermo Fisher Scientific, Rochester, NY) were added for a 16-hour incubation. Afterwards, medium containing viral particles was replaced with two milliliters of fresh DMEM supplemented with 10% FBS. To select transduced cells, puromycin was added to a final concentration of 10 µg/ml in cell culture medium. To induce the expression of shRNA and GFP, 500 ng/ml doxycycline (Fisher Scientific, Pittsburgh, PA) was added for 72 hours, and the percentage of GFP+ CAFs or RFP+ melanoma cells was determined using a Cytation 1 Cell Imaging Multi-Mode Reader (BioTek Instruments, Winooski, VT). The efficiency of β-catenin ablation by shRNA in CAFs was determined by Western blotting.

### siRNA transfection

Silencer® siRNAs targeting BRAF (siRNA ID:507), CRAF (siRNA ID: 1548), POSTN (siRNA ID: 20888) and scramble Silencer® siRNA control were purchased from Thermo Fisher Scientific (Rochester, NY). Briefly, CAFs were first seeded in 6-cm culture dishes to reach about 60-70% confluence. Subsequently, 25 pmol of siRNA and 7.5 µl of Lipofectamine® RNAiMAX (Thermo Fisher Scientific, Rochester, NY) in 500 µl of Opti-MEM I Reduced Serum Medium (Thermo Fisher Scientific, Rochester, NY) were added for a 24-hour incubation according to standard protocol. Next day, culture medium containing siRNA and RNAiMAX was replaced with regular DMEM for cell culture.

### MTT cell viability assay

In a 96-well plate, 5,000 CAFs were seeded per well in triplicate with 500 ng/ml doxycycline for a 72-hour induction. 200 µl of 0.5 mg/ml MTT solution (3-(4,5-dimethylthiazol-2-yl)-2,5-diphenyltetrazolium bromide) were added to each well and incubated for four hours at 37°C. After incubation, MTT solution was replaced with 200 µl of dimethyl sulfoxide (DMSO). The plate was incubated at room temperature on a rocking shaker with gentle shaking for five minutes before being transferred to a microplate reader for measuring the absorbance at 594 nm.

### Collagen gel contraction assay

Collagen gels were prepared as described previously^73^. Briefly, 3 mg/ml of type I collagen (Thermo Fisher Scientific, Rochester, NY), DMEM medium, FBS, and 1 N NaOH solution were mixed on ice at a volume ratio of 1:1.65:0.3:0.05 to make a 1 mg/ml collagen solution. 50,000 CAFs after a -72-hour doxycycline induction were then resuspended in 500 µl of 1 mg/ml collagen solution and transferred into one well of a 24-well tissue culture plate. After a 30-minute incubation in a humidified incubator at 37°C with 5% CO_2_, one ml of fresh medium was added on top of the gel for a 72-hour incubation. Afterwards, the gels were detached from the wall of each well and allowed for contraction as indicated. Pictures of the gels were taken using a Nikon digital camera every 24 hours, and ZEN 2.3 software was used to measure the diameters. The gel contraction percentage was quantified by dividing the difference of gel diameters at 0 hour and the 72th hours by the gel diameter at 0 hour.

### Confocal reflection microscopy

The experimental procedure was done in the same way as described for collagen gel contraction assay. After a 72-hour incubation, collagen fiber distribution was assessed using a Zeiss LSM 710 confocal microscope at 40X magnification. Images were acquired at a depth of at least 100 mm inside the gel to avoid edge effects. Images of at least ten areas of each gel were randomly captured for 3D reconstruction of the matrix using ImageJ (https://imagej.nih.gov). Fiber connectivity was calculated using ImageJ with the BoneJ plugin (http://bonej.org)74.

### *In vitro* scratch assay

CAFs were treated with 500 ng/ml doxycycline for 72 hours and then seeded in a 24-well tissue culture plate. When the cells reached 90% confluence, culture medium was replaced with serum-reduced medium (DMEM supplemented with 0.5% FBS) supplemented with 0.02 mg/ml mitomycin C (Sigma, St. Louis, MO), which inhibits cell proliferation. The fibroblast layer was scratched using a p200 pipet tip in the center of each well to form a rectangular cell-free area with a width of around 800 to 1000 µm. The cells were allowed to migrate for 48 hours, and then bright-field images were captured using a Leica DMi8 wide-field microscope. At least ten images were taken per well at each time point to calculate the width of scratched area using LAS X software (Leica Microsystems). The cell migration percentage was calculated using the following formula: (average width of scratched area at 48 hours average width of scratched area at 0 hour)/average width of scratched area at 0 hour.

### Atomic force microscopy

Collagen gels embedded with CAFs were generated as described in collagen gel contraction assay. After a three-day incubation, the gels were transferred to a petri dish and immersed in DMEM with 10% FBS. The stiffness of the gels was then measured using an atomic force microscope (NanoWizard 4a ver. 6.1.99, JPK Instruments) with a silicon nitride cantilever (CSC37, k = 0.3 N/m, MikroMasch). The Young’s modulus was calculated from the force-distance curves using a modified Hertz model of JPK Data Processing (version 6.1.99, JPK Instruments).

### 3D cell co-culture system

To form a 3D co-culture cell spheroid, 5,000 RFP-tagged A375 or SK-MEL-24 cells were mixed with 5,000 GFP-tagged CAFs in one well of a U-bottomed 96-well plate (Thermo Fisher Scientific, Rochester, NY). Spheroid was formed by centrifuging the plate at 1,000 g for 10 minutes followed by incubation at 37°C in a humidified incubator with 5% CO_2_ overnight. For drug resistance assay, spheroids were cultured using normal culture medium or medium containing 500 nM PLX4032 (Selleck Chemicals, Houston, TX) or 100 nM GDC0973 (Selleck Chemicals, Houston, TX) for 72 hours before they were collected for cell number counting. To count RFP+ melanoma cell number in the spheroids, 12 spheroids from each group were collected and digested using 2 mg/ml collagenase IV (Therma Fisher Scientific, MA) for one hour with stirring to generate a single cell suspension. A Countess II Automated Cell Counter (Thermo Fisher Scientific, Rochester, NY) was used to quantify melanoma cell number based on red fluorescence. Average melanoma cell number in each spheroid was calculated by dividing the total cell number by 12.

### Recombinant POSTN assay

For 2D recombinant POSTN assay, A375 cells were seeded in the wells of non-treated 6-well-plate (ThermoFisher, Waltham, MA) coated with POSTN (R&D Systems, Minneapolis, MN) and type I collagen. A375 cells cultured on cell culture plate coated only with type I collagen were used as a control. To coat the cell culture plate, one ml solution of 250 ng/ml type I collagen with or without 250 ng/ml POSTN (R&D Systems, Minneapolis, MN) was added to each well followed by one-hour incubation at room temperature. Afterwards, the solution was removed, and the plate was rinsed with PBS for three times. The cells were then seeded and cultured with or without 500 nM PLX4032 (Selleck Chemicals, Houston, TX) for 72 hours before they were collected for cell counting. Images of 2D-cultured RFP+ melanoma cells were captured using a Cytation 1 Cell Imaging Multi-Mode Reader (BioTek Instruments, Winooski, VT).

For 3D recombinant POSTN assay, 8,000 melanoma cells were seeded into one well of a U-bottomed 96-well plate (Thermo Fisher Scientific, Rochester, NY) to generate the spheroid as described above. The spheroids were cultured using normal culture medium or medium containing 500 nM PLX4032 (Selleck Chemicals, Houston, TX) or 100 nM GDC0973 (Selleck Chemicals, Houston, TX) with or without 250 ng/ml POSTN (R&D Systems, Minneapolis, MN) as indicated for 72 hours before they were collected for cell number counting. Images of the spheroids were captured using a Cytation 1 Cell Imaging Multi-Mode Reader (BioTek Instruments, Winooski, VT).

### Immunofluorescence staining and immunohistochemistry

For the cells cultured in the wells of the 8-well Nunc™ Lab-Tek™ II chamber slide (Thermo Fisher Scientific, Rochester, NY), the cells were fixed in 4% paraformaldehyde for 15 minutes at room temperature and then permeabilized using 0.1% Triton-100 for 10 minutes on ice. The cells were then washed three times with PBS and blocked using 10% BSA for one hour. Primary antibodies recognizing β-catenin (Sigma, MA1-301, 1:1000), α-SMA (Thermo Fisher, 14-9760-82, 1:200), FAP (Thermo Fisher, PA5-51057, 1:200), FSP-1 (Abcam, ab124805, 1:200), paxillin (BD Biosciences, 610051, 1:200), p-MLC2 (Cell Signaling Technology, 3671, 1:100), F-actin (Thermo Fisher, R415, 1:20), and Ki67 (BD Biosciences, 550609, 1:200) were then added to specific chambers and incubated overnight at 4°C. Next day, after washing with PBS for three times, an Alexa Fluor 488 or 594-conjugated secondary antibody (Invitrogen, Carlsbad, CA) was added for incubation to visualize the expression of proteins. Images were taken using a Zeiss LSM 710 confocal microscope and analyzed using ImageJ (NIH).

To quantify the fluorescent intensity of immunostained fibroblasts in chamber slides, the slides were photographed using a Cytation 1 multi-mode cell imager, and the fluorescence intensity in each cell was quantified using GEN 5.3 software (BioTek, Winooski, VT). In addition, the fluorescence intensity of nuclear β-catenin staining in each cell was quantified only in the nuclear area indicated by DAPI staining. The percentage of Ki67+ cells were quantified by dividing the number of Ki67+ cells by the total cell number in each field.

For co-cultured spheroids and mouse tumors, the spheroids and tumor tissues were washed once in PBS, fixed overnight in 4% paraformaldehyde, and then embedded in agarose gel for the preparation of paraffin blocks. Paraffin sections were cut at a thickness of 5 µm. Briefly, the sections were deparaffinized, rehydrated and then unmasked in citrate buffer (pH 6.0) using a microwave heating method. After washing with PBS, sections were blocked in 10% bovine serum albumin (BSA) (Thermo Fisher Scientific, Rochester, NY) in PBS and then incubated at 4°C overnight with indicated primary antibody. The following primary antibodies were used: TE-7 (fibroblast marker, Sigma Aldrich, CBL271, 1:200), α-SMA (CAF marker, Thermo Fisher, 14-9760-82, 1:200), GP-100 (melanoma marker, Abcam, sb137078, 1:50), Ki67 (BD Biosciences, 550609, 1:200), p-ERK (Cell Signaling Technology, 4370, 1:200), cleaved caspase-3 (Cell Signaling Technology, 9661, 1:200), and POSTN (Invitrogen, PA5-34641, 1:50). Next day, the slides were washed three times using PBS, incubated with the corresponding biotin-conjugated secondary antibodies (Vector Laboratories, Burlingame, CA) at room temperature for one hour, and then incubated with fluorochrome-labeled streptavidin (Vector Laboratories, Burlingame, CA) for another hour. The slides were mounted using VECTASHIELD Antifade Mounting Medium (Vector Laboratories, Burlingame, CA) and coverslipped. Images were acquired using a Nikon Eclipse 80i fluorescence microscope and quantified using ImageJ.

The numbers of CAFs in immunostained mouse (GP-100-) and human melanoma tissue sections (α-SMA or TE7+) were counted in each high-power field (40X) using the particle analysis function of ImageJ software (NIH). 4.5 high-power microscopic fields (40X) are equal to one square millimeter. Therefore, cell number per square millimeter was calculated by multiplying cell number counted in each field by 4.5. For each melanoma samples, at least 25 randomly selected fields are counted. A minimum of three independent experiments were done for each immunostaining quantification.

### TUNEL assay

TUENL assay was performed following the standard protocol provided by the manufacturer (Sigma, St. Louis, MO). Briefly, the cells cultured in the wells of the 8-well Nunc™ Lab-Tek™ II chamber slide (Thermo Fisher Scientific, Rochester, NY) were fixed in 4% paraformaldehyde for 15 minutes at room temperature and then permeabilized using 0.1% Triton-100 for 10 minutes on ice. The cells were then washed three times with PBS and blocked using 10% BSA for one hour. TUNEL reaction mixture was made by mixing the label solution and enzyme solution at a ratio of 9:1. 50 µl of TUNEL reaction mixture was added in each well, and the slide was incubated in a humidified atmosphere for 60 min at 37°C in the dark. Images were acquired using a Nikon Eclipse 80i fluorescence microscope and quantified using ImageJ. The percentage of TUNEL+ cells were quantified by dividing the number of TUNEL+ cells by the total cell number in each field.

### Tumor xenografting experiments

All animal experiments were performed in accordance with the protocol approved by the Institutional Animal Care and Use Committee (IACUC) of the University of Cincinnati (10-09-23-01). Human melanoma xenograft model was established using a combination of BRAF-mutant human melanoma cell lines, including A375 and SK-Mel-24, and CAF lines, including M27 and M50. To induce melanomas in mice, a mix of 1 × 10^6^ A375 cells and 1 × 10^6^ uninduced GFP/M50 or bcat-GFP/M50; or 1 × 10^6^ SK-Mel-24 cells and 1 × 10^6^ uninduced GFP/M27 or bcat-GFP/M27 in 50 µl of growth-factor-reduced Matrigel/PBS (1:1) (BD Biosciences) was injected intradermally into the flanks of 4-6-week-old NOD.Cg-Prkdc^scid^/J mice (Jackson Laboratory, stock number 001303), respectively. The mice were observed daily for tumor appearance. To induce β-catenin ablation, all mice were fed with doxycycline diet for four weeks after the tumors reached a volume size of approximately 50 cubic millimeters. The mice in each group received either saline buffer or drug treatment as indicated (vemurafenib/PLX4032 [Selleck Chemicals, Houston, TX], 50 mg/kg every other day; cobimetinib/GDC0973 [Selleck Chemicals, Houston, TX], 20mg/kg every other day). Meanwhile, the size of tumors was measured and recorded every other day until the end point when the tumor size exceeded 20% of body size. After the mice were euthanized, the tumors were harvested for various analyses.

### Collagen staining

Picrosirius red staining was performed according to the manufacturer’s instructions (American MasterTech, Lodi, CA). Briefly, paraffin sections of mouse melanomas were dewaxed, hydrated and stained in picrosirius red dye for one hour. After staining, the slides were washed with 1% acetic acid for one minute, and excessive buffer was removed from the slides by vigorous shaking. Sections were then dehydrated in 100% ethanol, cleared using xylene and mounted with permanent mounting medium (Vector lab, Burlingame, CA).

### Western blotting

Western blotting was performed using the standard protocol published elsewhere. The following primary antibodies were used: β-catenin (Sigma, MA1-301, 1:2000), p-β-catenin (Cell Signaling Technology, 9561, 1:500), β-actin (Cell Signaling Technology, 3700S, 1:2000), ERK1/2 (Cell Signaling Technology, 9107, 1:2000), p-ERK1/2 (Cell Signaling Technology, 4370, 1:2000), periostin (Invitrogen, PA5-34641, 1:500), p-GSK-3β (Cell Signaling Technology, 5558, 1:2000), GSK-3β (Cell Signaling Technology, 12456, 1:2000), BRAF (Cell Signaling Technology, 9433, 1:500), CRAF (Cell Signaling Technology, 9422, 1:500), Akt1 (Abcam, ab32505, 1:1000), p-Akt1 (Abcam, ab66138, 1:1000), and Lamin B (Invitrogen, 50-554-010, 1:1000). Membranes were incubated with primary antibodies overnight at 4°C, washed three times using TBST (Tris-buffered saline, 0.1% Tween 20), and then incubated with IRDye 800CW donkey anti-rabbit IgG secondary antibody (LI-COR, 926-32213, 1:2000) or IRDye 680RD donkey anti-mouse IgG secondary antibody (LI-COR, 926-68072, 1:2000). The membranes were then washed again using TBST and processed for scanning and visualization using an Odyssey CLx imaging system (LI-COR, model #9140). Protein bands were quantified using ImageJ as published before^75^.

### Co-immunoprecipitation

Coimmunoprecipitation was performed using a Pierce Crosslink Immunoprecipitation Kit according to the manufacturer’s instruction (ThermoFisher, Waltham, MA). Briefly, 10 mg of BRAF antibody (Cell Signaling Technology, 9433) or mouse IgG (ThermoFisher, 31903) were added to a spin column containing 20 ml of resin slurry in a 2-ml collection tube. The mix was incubated at room temperature with gentle rotation for one hour. After incubation, the spin columns were centrifuged, then the antibody-bound resin was rinsed with 1X coupling buffer three times to remove unbound antibodies. Fifty µl of 2.5 mM disuccinimidyl suberate (DSS) crosslinker solution was added to crosslink the antibody to the resin. Subsequently, 750 mg of protein extract of M50 or PLX4032-treated M50 was added to the spin column and rotated end-over-end at 4°C overnight. After incubation, the column was washed three times with 1X TBS and one time with 1X conditioning buffer. Fifty ml of elution buffer was used to elute the samples from the column. The eluate was boiled with sample buffer at 95°C for five minutes for Western blotting.

### Chromatin immunoprecipitation

ChIP assay was performed according to the manufacturer’s instruction (Pierce™ Agarose ChIP Kit, Invitrogen, 26156). Briefly, two 10-cm dishes of confluent CAFs were collected for each ChIP assay (1 × 10^7^ -5 × 10^7^ cells). Crosslinking, cell pellet isolation, cell lysis, and MNase digestion were performed as exactly described in the instruction. The digested chromatin was collected by centrifugating at 9000 x g for five minutes for immunoprecipitation.

500 µl diluted lysate was incubated with the antibody against β-catenin (10 µg; Sigma, MA1-301, 1:2000) overnight at 4 °C with gentle rocking. ChIP assay using normal rabbit IgG (Invitrogen, 10500C) was performed as mock control. 20 µl of ChIP Grade Protein A/G Plus Agarose was added to each IP and incubated for one hour at 4 °C. After several washing steps, IP elution was performed by adding 150 µl 1X IP elution buffer to the washed resin in the collection tube and incubated at 65°C for 30 minutes with shaking. Six µl of 5M NaCl and 2 µl of 20 mg/ml proteinase K were added to the tube. The tube was then centrifuged at 6000 x g for one minute to stop the reaction. All IP samples were placed in a 65 °C heat block for one and a half hours. For DNA recovery, 50 µl of DNA elution solution were used to elute DNA.

Immunoprecipitated DNA fragment were subjected to quantitative real-time PCR with primer sets that amplify POSTN gene fragment (the promoter region) containing potential LEF/TCF-binding sites (Forward: CAGCCTTTACCCCCTTGTGA Reverse: GAACAATTGACTCACTGCATGTT). The primer pair amplifying the intron 1 region of Axin2 gene that contains previously reported LEF/TCF-binding sites was used as a positive control (Forward: TTTTGCATAAAAGGGGCGGC, Reverse: CGGCACTCACCATTGATCCC)^76^. ChIP-qPCR data were normalized using the percent input method^77^.

### RNA-Seq

Braf^V600E^; Pten^lox5/lox5^ melanomas carrying either wildtype Fb or β-catenin-deficient fibroblasts (bcat/Fb) were generated as we published previously. Control Fbs or bcat/Fbs were isolated from respective melanoma by flow cytometer cell sorting (FACS) using PDGFR-α as a marker. Normal fibroblasts were isolated from B6 mouse skin. RNA was extracted from normal fibroblasts, control Fbs or bcat/Fbs for RNA-Seq using Illumina NextSeq 550 system. The data were examined and analyzed using A.I.R. (Sequentia Biotech, Barcelona, Spain) and filtered according to the fold difference in expression. The Venn diagram was generated using Venny 2.1.0 (https://bioinfogp.cnb.csic.es/tools/venny/). The functional annotation clustering analysis was performed using DAVID Bioinformatics Resources 6.8^78^.

### Quantitative Real-Time PCR

RNA was extracted using PureLink™ RNA Mini Kit following the standard protocol (ThermoFisher, Waltham, MA). RNA concentration was determined using a NanoDrop™ spectrophotometer (ThermoFisher, Waltham, MA). First-strand cDNAs were synthesized for real-time PCR by reverse transcription using the SuperScript^TM^ IV first-strand synthesis system (ThermoFisher, Waltham, MA). qPCR primers for POSTN were purchased from realtimeprimers.com (Philadelphia, PA), and qPCR reactions were performed using power SYBR green master mix on a StepOnePlus real-time PCR system (Applied Biosystems). Relative gene expression levels were normalized to GAPDH.

### Statistical analysis

All quantitative results were obtained from a minimum of three independent experiments. All data were analyzed using GraphPad Prism 8 software package (GraphPad Software Inc., San Diego, CA) and expressed as the mean ± SEM. Differences between means were determined by Student’s t-tests and were considered statistically significant at P < 0.05.

## Supporting information

DNA sequence for periostin promoter region

## ACKNOWLEDGMENTS

This work was supported by the Cincinnati Cancer Center - Mentor-Mentee Award (YZ), the Harry J Lloyd Trust Research Award (YZ) and the CCTST Pilot Translational Research & Innovative Core Grant (YZ). We thank the University of Cincinnati Biorepository and the Melanoma Biorepository at the University of Colorado Anschutz Medical Campus for providing human melanoma tissue sections. We thank Drs. Kentaro Iwasawa and Takanori Takebe at the Cincinnati Children’s Hospital Medical Center for helping with AFM. We thank Ryan Dickerson from the Educational Technology Division at the University of Central Florida for assisting with illustration in Fig 10.

## COMPETING INTERESTS

The authors declare no competing interests.

## CONTRIBUTIONS

T.L. and Y.Z. designed the experiments; T.L. Y.X. and L.Z. performed the experiments and collected the data. Y.Z., T.L., and L.Z. analyzed the data; Y.Z., T.A., and T.L. wrote the main manuscript text, and T.L. and Y.Z. prepared the figures. All authors reviewed and approved the final version to be published. All authors agree to be accountable for all aspects of the work and ensure that questions related to the accuracy or integrity of any part of the work are appropriately investigated and resolved; further, the authors have confidence in the integrity of the contributions of their coauthors.

**Correspondence** and requests for materials should be addressed to Y.Z.

---

**Fig. S1.**
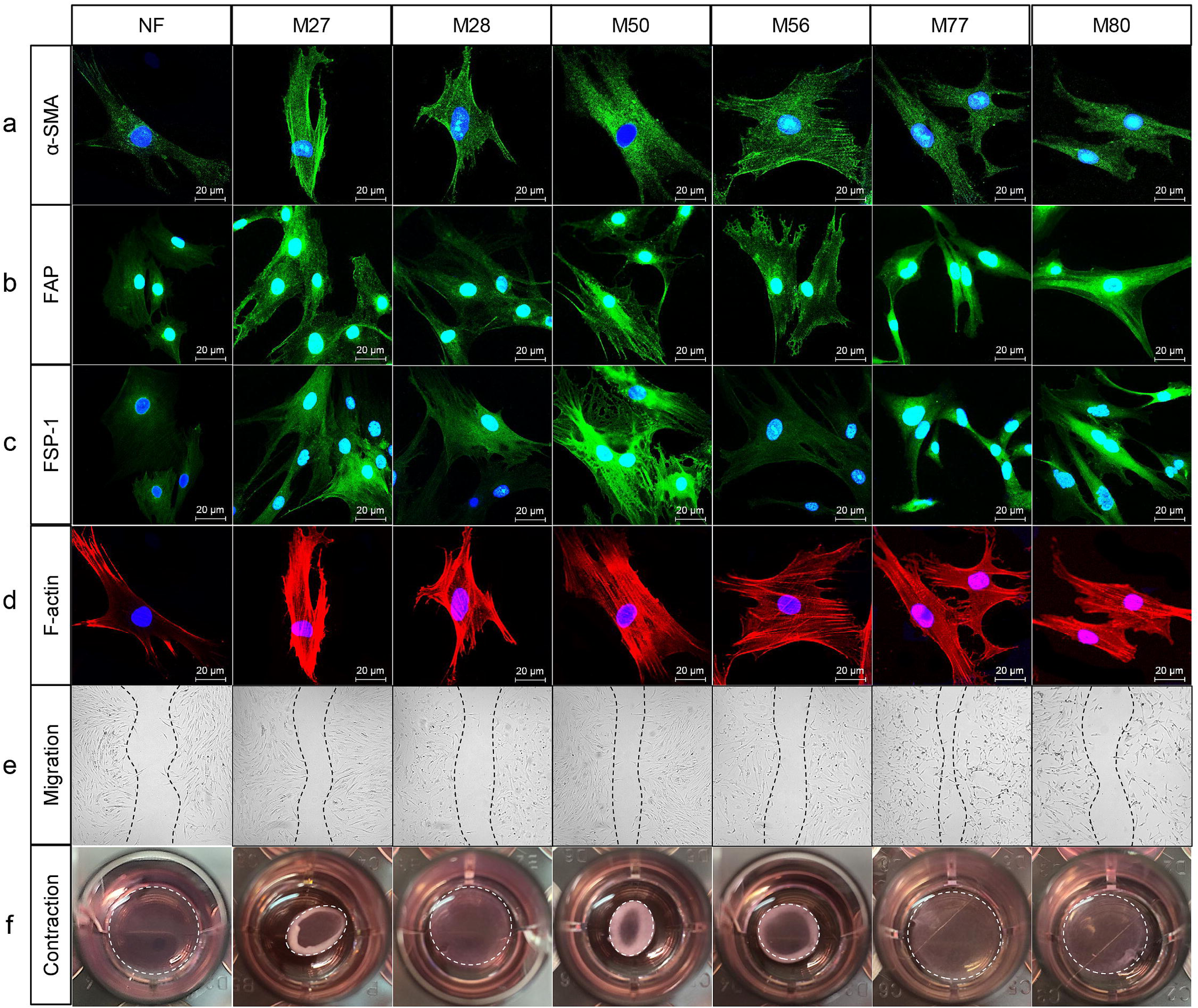
Characterization of CAFs isolated from surgically excised human melanoma tissues. **a-d**. Representative images show α-SMA, FAP, FSP1, and F-actin staining in NF isolated from normal human skin and M27, M28, M50, M56, M77, and M80 isolated from human melanoma tissues. Scale bar: 20 μ **e**. The scratch assay was used to evaluate the migratory abilities of the fibroblasts. Representative images were taken after 48 hours. n=3. **f**. Collagen gel contraction assay was performed to assess the ECM remodeling abilities of the fibroblasts. White circles indicate the size of the gel after a 72-hour contraction. n=3.

**Fig. S2.**
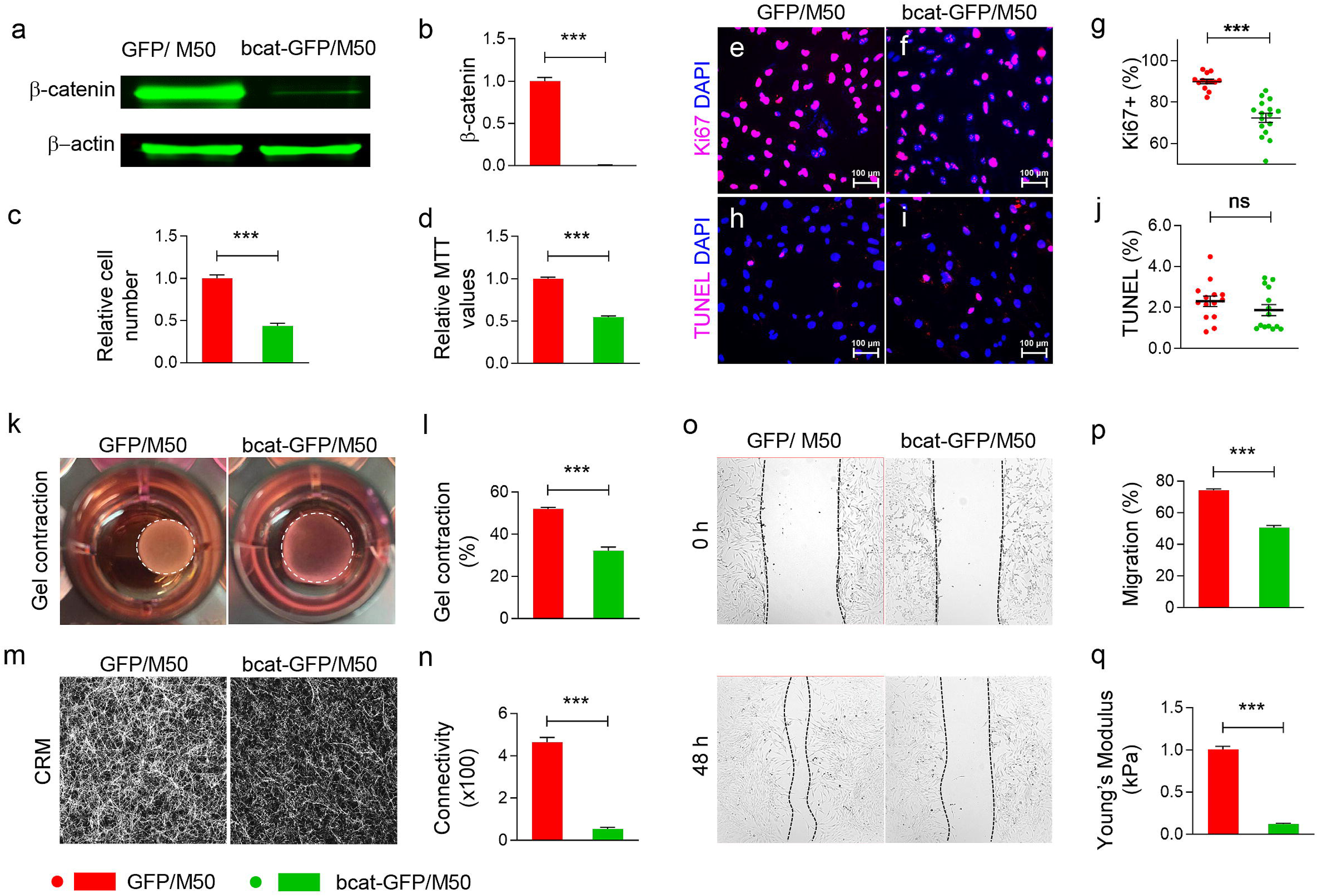
β-catenin is essential for the biological properties of CAFs. **a**. Western blotting of β-catenin expression in GFP/M50 and bcat-GFP/M50 after doxycycline induction. **b.** Bar graph shows fluorescence intensities of β-catenin bands of GFP/M50 and bcat-GFP/M50. The intensities of β-catenin bands were normalized to the corresponding β-actin bands. n=3. **c**. Relative numbers of GFP/M50 and bcat-GFP/M50 after a 72-hours doxycycline induction. n=3. **d.** Bar graph shows relative MTT values from GFP/M50 and bcat-GFP/M50 after a 72-hours doxycycline induction. n=3. **e-f**. Images show Ki67 staining of GFP/M50 and bcat-GFP/M50 with DAPI nuclear counterstaining (blue). Scale bar: 100 μm. **g**. Scatter graph shows the percentages of Ki67+ cells in DAPI+ cells in GFP/M50 and bcat-GFP/M50. Each data point represents the Ki67+ percentage in each 20X field. At least 13 fields were photographed and quantified. **h-i**. Images show TUNEL staining of GFP/M50 and bcat-GFP/M50 with DAPI nuclear counterstaining (blue). Scale bar: 100 μm. **j**. Scatter graph shows the percentages of TUNEL+ cells in DAPI+ cells in GFP/M50 and bcat-GFP/M50. Each data point represents the TUNEL+ percentage in each 20X field. At least 14 fields were photographed and quantified. **k**. Representative images of collagen gels contracted by GFP/M50 and bcat-GFP/M50 after 72 hours. **l**. Bar graph shows percentages of gel contraction from three independent experiments. **m**. Representative CRM images of the gels embedded with GFP/M50 and bcat-GFP/M50 after 72 hours. **n**. The graph shows the connectivity of collagen fibers in gels remodeled by GFP/M50 and bcat-GFP/M50 from three independent experiments. **o**. Images show GFP/M50 and bcat-GFP/M50 migration at 0 and the 48th hour in an *in vitro* scratch assay. Black short dash lines depict the frontlines of GFP/M50 and bcat-GFP/M50 at indicated time points. **p**. Bar graph shows migration percentage of GFP/M50 and bcat-GFP/M50 at the 48th hour. n=4. **q**. The graph shows Young’s modulus of collagen gels, which were remodeled by GFP/M50 and bcat-GFP/M50 for 72 hours, measured by AFM. n≥26. For all graphs, data are quantified and shown as mean ± SEM. *** *P* ≤ 0.001. ns: not significant.

**Fig. S3.**
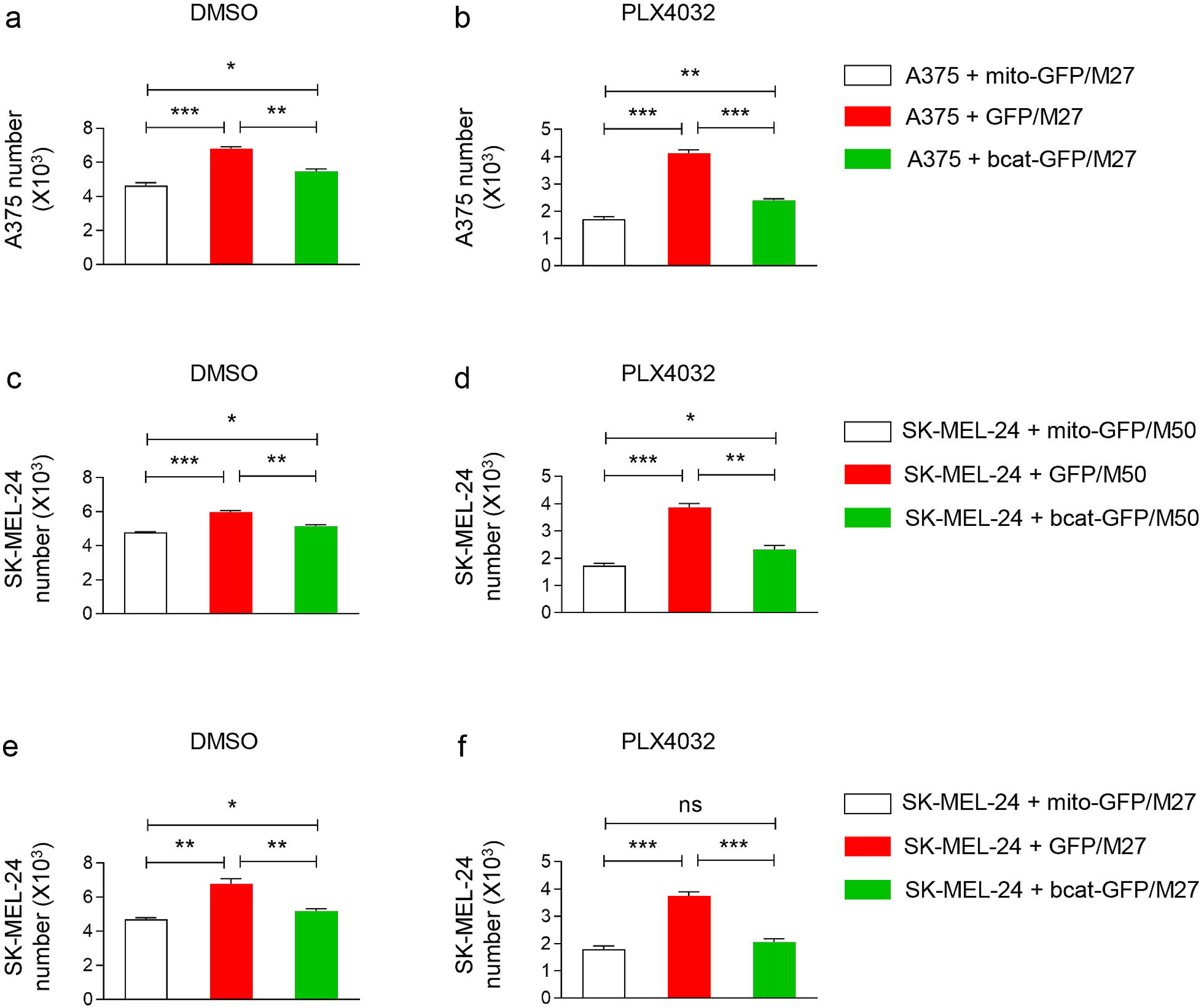
β-catenin plays an important role in the contribution of CAFs to melanoma resistance to BRAFi. **a-b**. Bar graph shows A375 numbers in 3D spheroids co-cultured with mito-GFP/M27, GFP/M27, or bcat-GFP/M27 and treated with DMSO or PLX4032. n=3. **c-d**. Bar graph shows SK-MEL-24 numbers in 3D spheroids co-cultured with mito-GFP/M50, GFP/M50, or bcat-GFP/M50 and treated with DMSO or PLX4032. n=3. **e-f**. Bar graph shows SK-MEL-24 numbers in 3D spheroids co-cultured with mito-GFP/M27, GFP/M27, or bcat-GFP/M27 and treated with DMSO or PLX4032. n=3. For all graphs, data are shown as mean ± SEM. * *P* ≤ 0.05. ** *P* ≤ 0.01; *** *P* ≤ 0.001. ns: not significant.

**Fig. S4.**
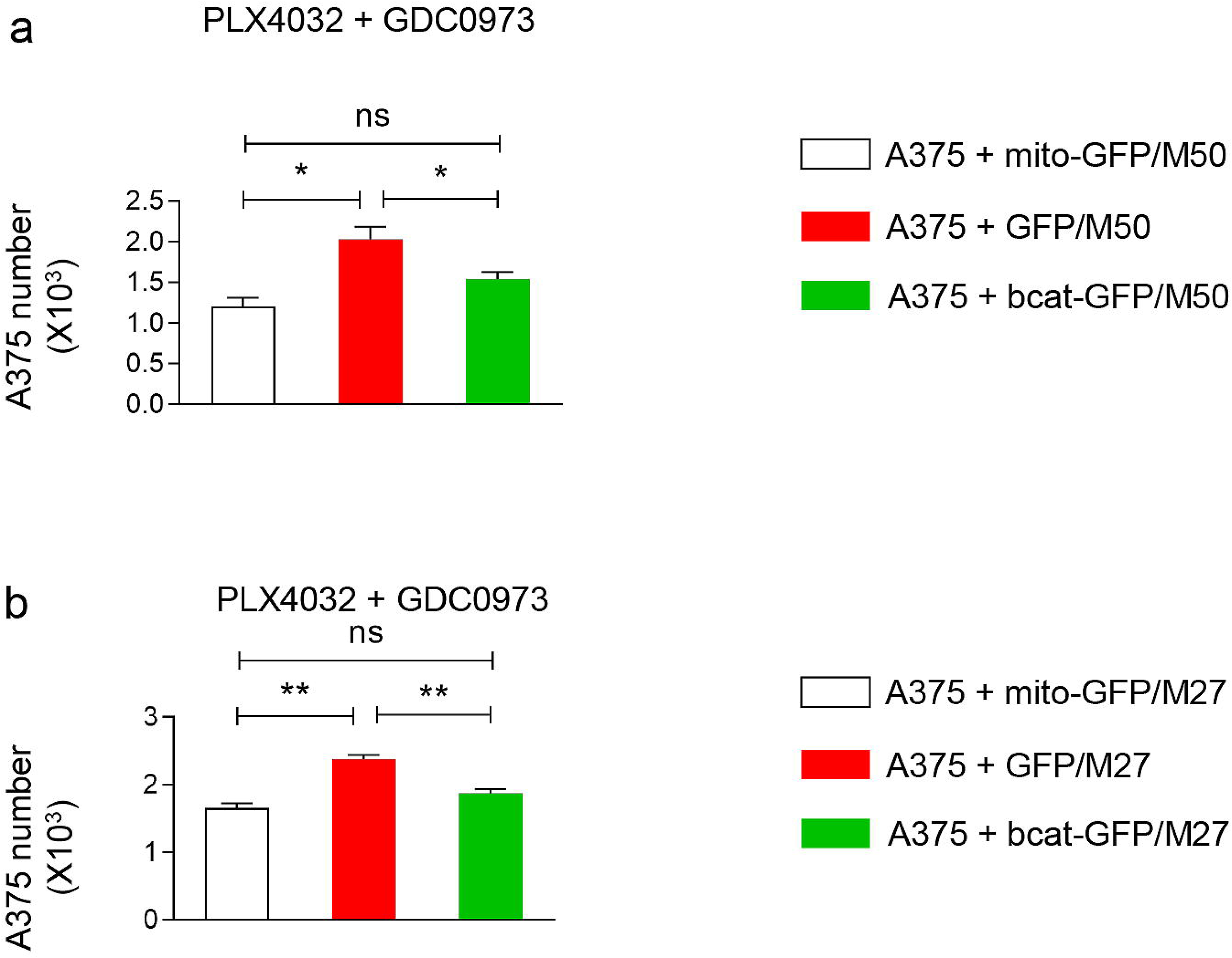
β-catenin plays an important role in the contribution of CAFs to melanoma resistance to BRAFi and MEKi. **a**. Bar graph shows A375 numbers in 3D spheroids co-cultured with mito-GFP/M50, GFP/M50, or bcat-GFP/M50 and treated with PLX4032 and GDCC0973. n=3. **b**. Bar graph shows A375 numbers in 3D spheroids co-cultured with mito-GFP/M27, GFP/M27, or bcat-GFP/M27 and treated with PLX4032 and GDC0973. n=3. For all graphs, data are shown as mean ± SEM. * *P* ≤ 0.05. ** *P* ≤ 0.01; *** *P* ≤ 0.001. ns: not significant.

**Fig. S5.**
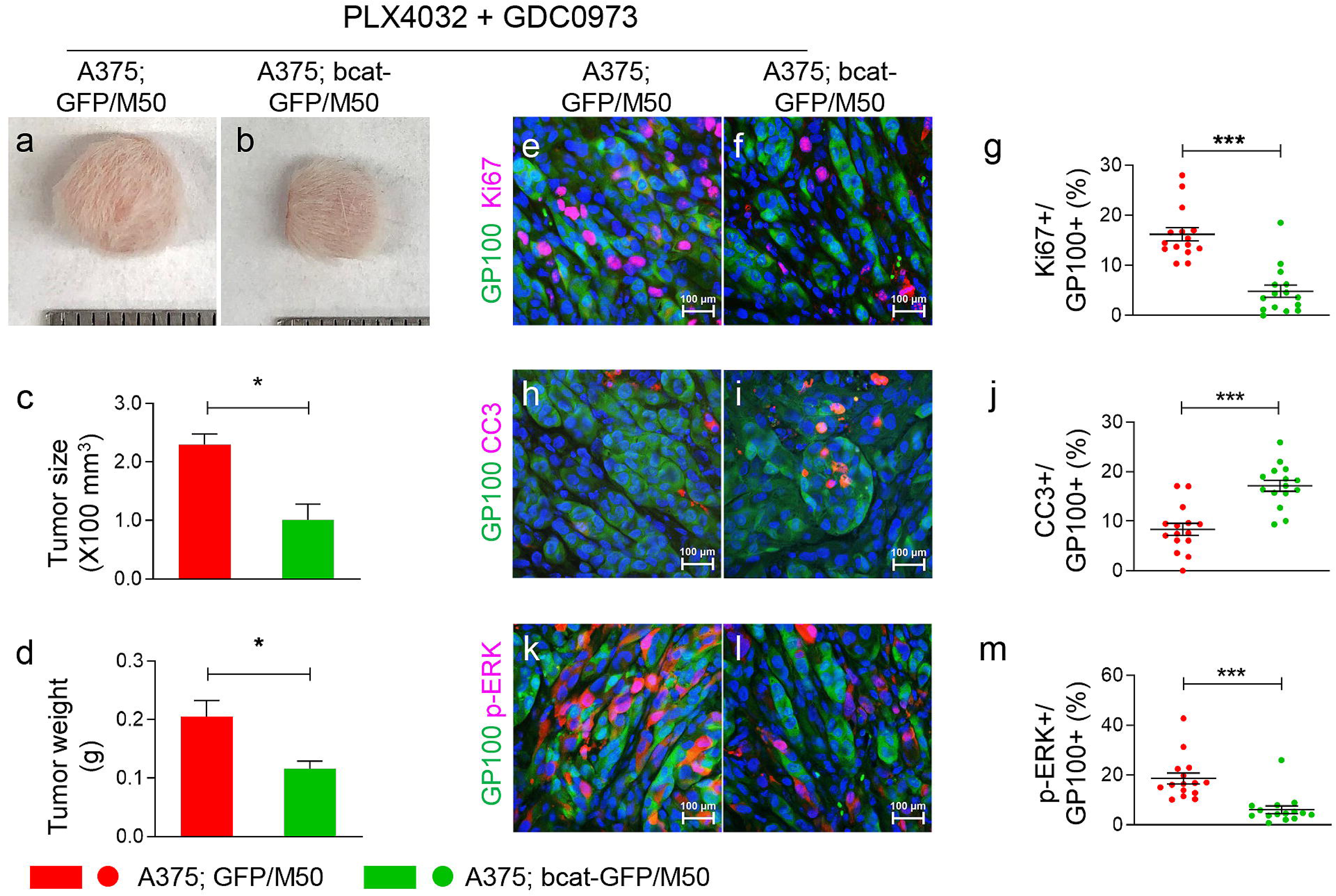
β-catenin ablation in CAFs sensitizes BRAF-mutant melanoma cells to BRAFi/MEKi *in vivo*. **a-b**. Representative pictures of melanoma xenografts in indicated groups 28 days after either saline or PLX4032 and GDC0973 treatment. **c-d**. Comparative analysis of the volumes (**c**) and weights (**d**) of melanoma xenografts between two experimental groups. n=3. **e-f**. Images show GP-100 (melanoma cells, green) and Ki67 (red) staining of melanoma tissues sections collected from each group. **g**. Scatter graph shows the percentages of GP100+ melanoma cells that are Ki67+in indicated groups. Each data point represents the percentage of Ki67+ melanoma cells in one 40X field. **h-i**. Images show GP-100 (green) and CC3+ (red) staining of melanoma tissues sections collected from each group. **j**. Scatter graph shows the percentages of GP100+ melanoma cells that are CC3+in indicated groups. Each data point represents the percentage of CC3+ melanoma cells in one 40X field. **k-l**. Images show GP-100 (green) and p-ERK (red) staining of melanoma tissues sections collected from each group. **m**. Scatter graph shows the percentages of GP100+ melanoma cells that are p-ERK+ in indicated groups. Each data point represents the percentage of p-ERK+ melanoma cells in one 40X field. For all staining pictures: n≥15. Scale bar: 100 µm. In all graphs, data are shown as mean ± SEM. * P ≤ 0.05; ** P ≤ 0.01; *** P ≤ 0.001. ns: not significant.

**Fig. S6.**
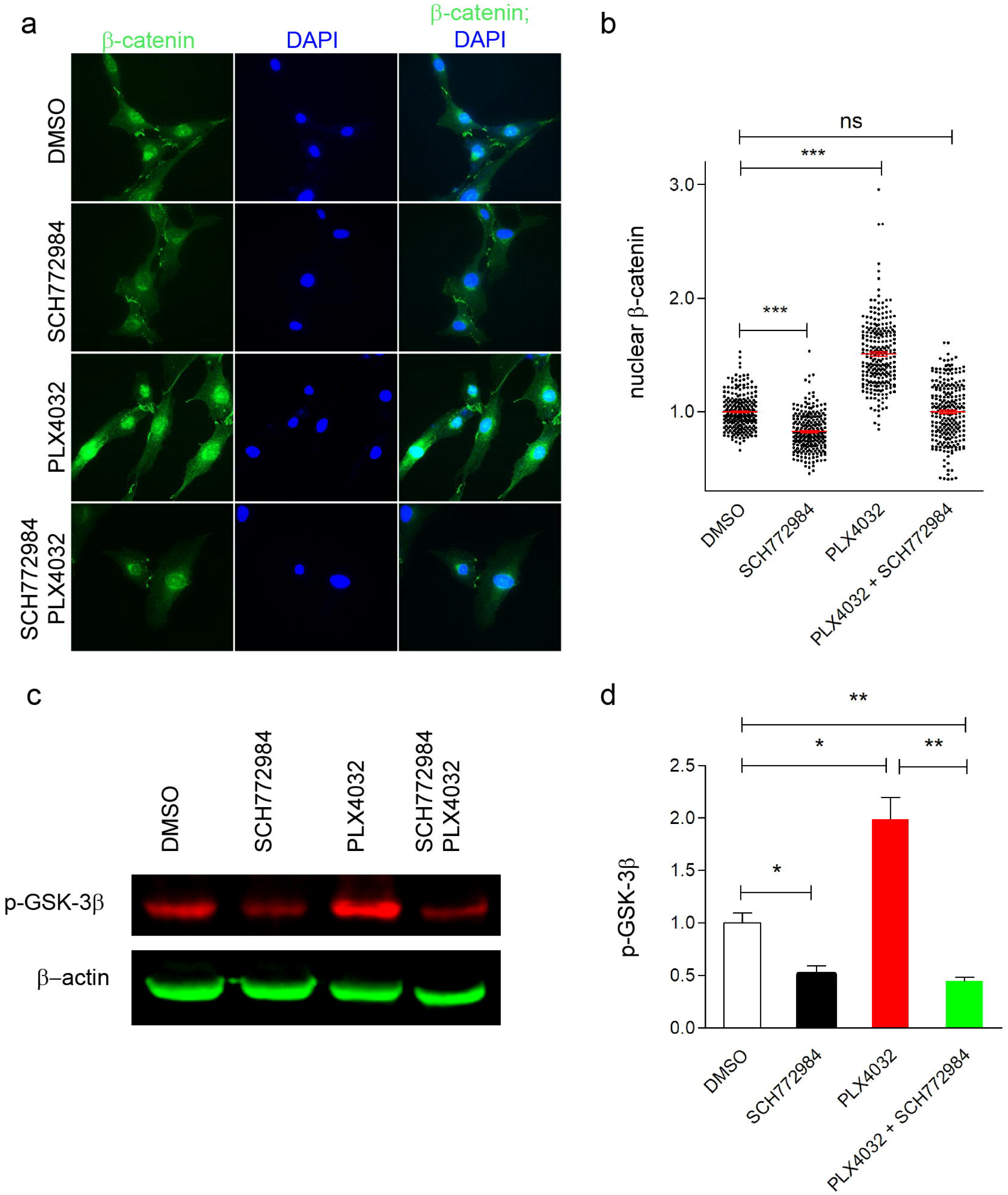
BRAF/CRAF dimerization activates ERK signaling to drive β-catenin nuclear translocation in CAFs. **a**. Representative images show β-catenin immunostaining of M50 treated with DMSO, SCH772984, PLX4032, or PLX4032/SCH772984. The nuclei were counterstained blue with DAPI. **b**. Scatter graph shows relative nuclear β-catenin expressions in M50 treated with DMSO, SCH772984, PLX4032, or PLX4032/SCH772984. The expression level of nuclear β-catenin was normalized by the mean nuclear β-catenin expression in M50 treated with DMSO for a 72-hour treatment. n≥100. **c**. Western blotting of p-GSK-3β expression in M50 treated by DMSO, SCH772984, PLX4032, or PLX4032/SCH772984 for two hours. **d**. Bar charts show quantification of fluorescence intensities of p-GSK-3β bands. n=3. Data are shown as mean ± SEM. * *P* ≤ 0.05. ** *P* ≤ 0.01; *** *P* ≤ 0.001. ns: not significant.

**Fig. S7.**
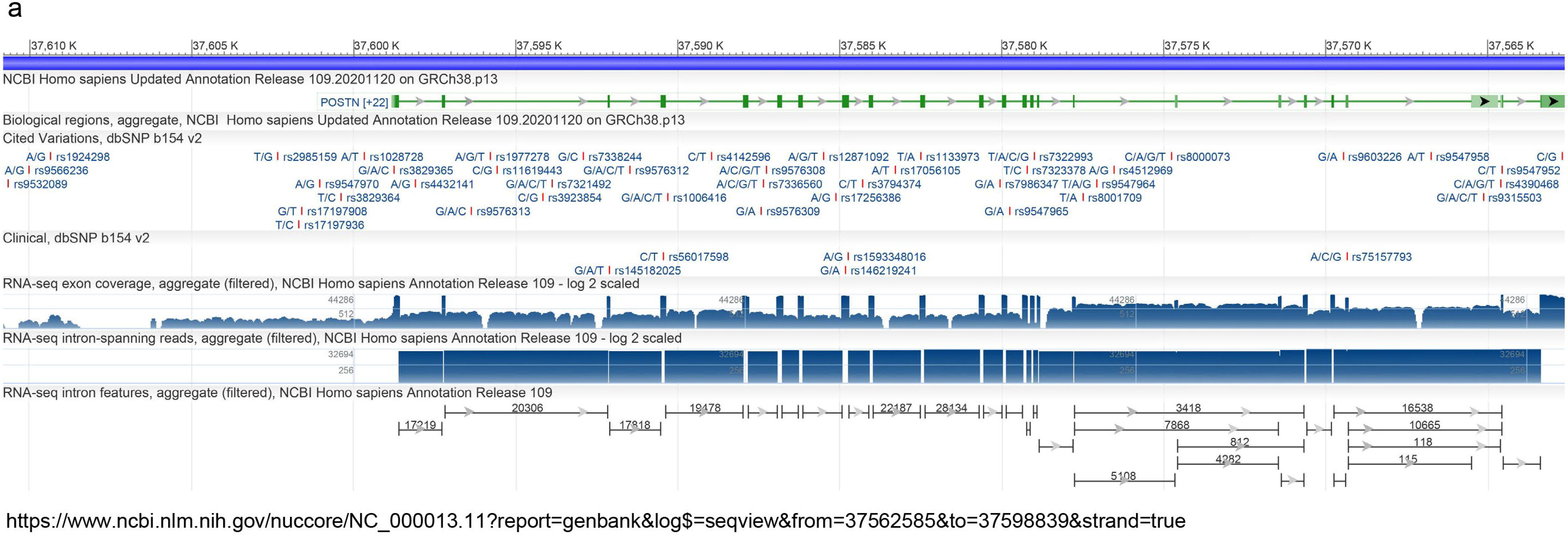
The putative human POSTN genomic locus (chr13:37562585-37598839, NCBI Homo sapiens updated annotation release 109.2020120 on GRCh38.p13).

## Notes

https://www.ncbi.nlm.nih.gov/geo/query/acc.cgi

### Competing Interest Statement

The authors have declared no competing interest.

