## Supplementary material for "BRAF inhibitors reprogram cancer-associated fibroblasts to drive matrix remodeling and therapeutic escape in melanoma": DNA sequence for periostin promoter region

```
Nucleotides in Exon 1 of POSTN are capitalized.
Blue marks the bases of the coding sequence in Exon 1.
Predicted TCF/LEF binding region(chr13:37599016-37599023)are colored green.

Primers are designed binding to sequences that are highlighted in yellow.


cttatttcat atcttacaca agttttaaat ctatccagag tttgttttta  37599240
atcaacagcc tttaccccct tgtgattgtc agactcgcat ctacctttgt  37599190
tttctggtaa aaataataat aataataatc tttcagttct gatgtgaact  37599140
gcaataacac ctaacaataa tcttgagcac acagacatta tacattctac  37599090
tctggaaagg attgcagaat atctcttaaa actcaacaaa agaatttttc  37599040
ttaaaaaccc tctaagATAC AAAGgaataa aactgagact taaacatgca  37598990
gtgagtcaat tgttcatatg attaaaaata agtaccttct ttataatgaa  37598940
aaggaaaagt agctcaatgt gttccttaaa tataactaac caaaacaaat  37598890
cttagctggc aatttgaagt tgccgatgct tcctggaaag agttcagact  37598840
CTCAGGTTGA TGCAGTGTTC CCTCCCACAA CTCTGACATG TATATAAATT  37598790
CTGAGCTCTC CAAAGCCCAC TGCCAGTTCT CTTCGGGGAC TAACTGCAAC  37598740
GGAGAGACTC AAGATGATTC CCTTTTTACC CATGTTTTCT CTACTATTGC  37598690
TGCTTATTGT TAACCCTATA AACGCCAACA ATCATTATGA CAAGATCTTG  37598640
GCTCATAGTC GTATCAGGGG TCGGGACCAA GGgtaagtga gtggtttggt  37598590
tttataaacc atttttcttt ttcctcagct tgttagatgt tcaagtatgc  37598540
ataaagtttc tcatatatgc ctcagggttt tttttcctaa ttattataaa  37598490
gtaataggaa aaaaaggaaa atgtgagtat tctgtgattt atctatgcac  37598440
ttttaagctt taagacttaa ttgctctcta aataccttga atataattgc  37598390
tttcttatga aatgttatat aattctaatc taattaacac aatttagctg  37598340
ggcattcttt tataattcca tttatttatg tttttatttt cttccaaaaa  37598290
```
